# Structural characterization of human RPA70N association with DNA damage response proteins

**DOI:** 10.1101/2022.07.14.500000

**Authors:** Yeyao Wu, Ning Zang, Wangmi Fu, Chun Zhou

**Author notes:** These authors contributed equally (Yeyao Wu, Ning Zang, Wangmi Fu).

## Abstract

The heterotrimeric Replication protein A (RPA) is the ubiquitous eukaryotic single-stranded DNA (ssDNA) binding protein and participates in nearly all aspects of DNA metabolism, especially DNA damage response. The N-terminal OB domain of the RPA70 subunit (RPA70N) is a major protein-protein interaction element for RPA. Previous crystallography studies of RPA70N with p53, DNA2 and PrimPol fragments revealed that RPA70N binds to amphipathic peptides that mimics ssDNA. NMR chemical-shift studies also provided valuable information of RPA70N residues interacting with target sequences. However, it is still not clear how RPA70N recognizes and distinguishes such a diverse group of proteins. Here we present high resolution crystal structures of RPA70N in complex with peptides from HelB, ATRIP, RMI1, WRN and BLM. The structures showed that in addition to the ssDNA mimicry mode of interaction, RPA70N employs multiple ways to bind its partners, some of which may serve to increase the avidity of RPA70N binding.

## Introduction

Replication protein A (RPA) is a heterotrimeric protein complex composed of the RPA70, RPA32 and RPA14 subunits (Fairman & Stillman, 1988; Wood *et al*, 1988). It is the major eukaryotic single-stranded DNA (ssDNA) binding protein and has an affinity for ssDNA in the range of 10^−9^-10^−10^ M (Blackwell & Borowiec, 1994; Iftode *et al*, 1999; Kim *et al*, 1994; Kim *et al*, 1992; Wold, 1997). Several oligonucleotide binding (OB) domains (RPA70A, 70B, 70C and RPA32D) form the core ssDNA binding region (Bochkarev *et al*, 1997; Bochkareva *et al*, 2002; Fan & Pavletich, 2012; Flynn & Zou, 2010; Murzin, 1993; Yates *et al*, 2018). Due to its high affinity for ssDNA, RPA is involved in almost all aspects of DNA replication, repair, and recombination(Caldwell & Spies, 2020; Chen & Wold, 2014; Fanning *et al*, 2006; Iftode *et al*., 1999; Marechal & Zou, 2015; Wold, 1997; Zou *et al*, 2006). It helps to protect ssDNA from nucleolytic degradation and prevents ssDNA entanglement by removing DNA secondary structures.

In addition to its ssDNA binding function, RPA also serves as a beacon to recruit a plethora of protein factors involved in DNA metabolism mostly through the RPA70N and RPA32 winged-helix domains(Awate & Brosh, 2017; Caldwell & Spies, 2020; Fanning *et al*., 2006; Marechal & Zou, 2015). The RPA70N domain adopts an OB fold with a five-stranded anti-parallel beta-barrel but has very weak ssDNA affinity (Jacobs *et al*, 1999). Its primary role is to mediate protein-protein interaction with its basic and hydrophobic cleft as first shown by a series of studies of RPA70N interacting with p53 (Abramova *et al*, 1997; Dutta *et al*, 1993; Li & Botchan, 1993). RPA70N binds to p53 transactivation domain to coordinate DNA repair with the p53-dependent checkpoint control. In the crystal structure of RPA70N-p53 complex, the acid-hydrophobic peptide of p53 is shown to interact with the complementary basic and hydrophobic groove, mimicking ssDNA binding to OB domains (Bochkareva *et al*, 2005). RPA70N also binds to the N-terminus of ATRIP and is responsible for recruiting the ATR-ATRIP complex to DNA damage sites to initiate cell-cycle checkpoint (Ball *et al*, 2005; Namiki & Zou, 2006; Zou & Elledge, 2003). In addition, RPA70N mediates the interaction of RPA with the MRN complex and the 9-1-1 complex to protect replication forks during DNA damage response, through binding to MRE11 and Rad9 (Oakley *et al*, 2009; Robison *et al*, 2004; Wu *et al*, 2005; Xu *et al*, 2008b). Besides cell cycle regulatory proteins, many helicases involved in DNA repair interact with RPA70N as well (Awate & Brosh, 2017). Both BLM (Sgs1) and DNA2 interact with RPA and form a complex to carry out long range DNA resection during double strand DNA break repair (Cejka *et al*, 2010; Gravel *et al*, 2008; Nimonkar *et al*, 2011; Zhu *et al*, 2008). RPA was proposed to recruit both DNA2 and BLM through RPA70N, stimulating the helicase activity of BLM while enhancing the nuclease activity of DNA2 by removing DNA secondary structures (Brosh *et al*, 2000; Doherty *et al*, 2005; Nimonkar *et al*., 2011; Zhou *et al*, 2015). WRN, HelB and FancJ also binds to RPA through RPA70N and the presence of RPA greatly enhanced their helicase activities (Brosh *et al*, 1999; Doherty *et al*., 2005; Guler *et al*, 2012; Gupta *et al*, 2007; Hormeno *et al*, 2022; Shen *et al*, 1998; Shen *et al*, 2003; Suhasini *et al*, 2009; Tkac *et al*, 2016). Moreover, many other proteins involved in DNA repair and replication interact with RPA70N too. For example, the RMI1 component of the BTR complex (BLM-Topo IIIα -RMI1-RMI2) and PrimPol (DNA primase and DNA polymerase) directly associate with RPA70N (Dornreiter *et al*, 1992; Guilliam *et al*, 2017; Shorrocks *et al*, 2021; Wan *et al*, 2013; Xue *et al*, 2013).

In general, most of the proteins interacting with RPA70N utilize a motif around 20 amino acids long with a mixture of acidic and hydrophobic residues (Shorrocks *et al*., 2021). For these motifs, the exact sequence doesn’t share much homology despite the similarity in overall composition, indicating each motif could bind to RPA70N differently. To better understand the mechanism of RPA70N-mediated target protein recruitment, we set out to determine complex structures of RPA70N with peptide motifs that bind to it. So far quite a few studies employed NMR chemical shifts to probe the interaction sites of RPA70N with partner proteins (Guler *et al*., 2012; Kang *et al*, 2018; Liu *et al*, 2011; Ning *et al*, 2015; Xu *et al*., 2008b; Yeom *et al*, 2019). While the NMR chemical shift information is useful in identifying potential residues involved in binding, due to the transient nature of the interactions, the complex structures were not resolved by NMR. Several crystal structures of RPA70N in complex with bound peptide were reported, namely RPA70N-p53, RPA70N-DNA2, RPA70N-PrimePol and Rfa1N-Ddc2 (Bochkareva *et al*., 2005; Deshpande *et al*, 2017; Guilliam *et al*., 2017; Zhou *et al*., 2015). This is mainly due to the weak affinity between RPA70N and the protein sequence it recognizes, crystallization attempts often yield crystals of RPA70N itself without peptide bound. To overcome this problem, we fused target sequence to the C-terminus of RPA70N with a flexible linker in between. By adjusting the linker length, we managed to crystalize and determine the structure of RPA70N in complex with HelB, ATRIP, RMI1, WRN and two BLM peptides (Supplemental Table S1-3).

## Results

### Structure of the RPA70N-HelB peptide complex

HelB is a conserved helicase involved in DNA replication initiation, replication stress responses and negatively regulates DNA end-resection (Guler *et al*., 2012; Hazeslip *et al*, 2020; Taneja *et al*, 2002; Tkac *et al*., 2016). It has an RPA binding motif located in the helicase domain and its recruitment to chromatin correlates with the level of replication protein A (Guler *et al*., 2012). We crystallized a human HelB helicase peptide (residues 496 to 519) with the human RPA70N (residues 1 to 120) using the fusion strategy in space group P41212 and there is one molecule in the asymmetry unit (Figure 1A and B, Supplemental Table S2). Crystal packing analysis shows that the HelB peptide from one fusion protein is bound by a neighboring RPA70N molecule (Supplemental Figure S1). The electron desnity of the fusion linker is not observed as it is highly flexible. In the 1.6 Å structure, residues 496 to 517 of HelB form a 4-turn helix followed by a β turn and a 3_10_ helix (Figure 1B). The curved β sheet of RPA70 and the extending L12, L45 loops form a shallow groove where the amphipathic helix of HelB sits (Figure 1B and C). The negatively charged residues E496, E499 and D506 of HelB formed hydrogen bond and salt bridges with R81, T60, Q61 and R41 (Figure 1D). The hydrophobic residues V500, C504 and F507 of HelB packs against a broad hydrophobic patch formed by I83, M97, I95, V93, L87 and M57 (Figure 1D). The mixed basic and hydrophobic character of the RPA70N groove complements the acidic-hydrophobic nature of the HelB peptide (Figure 1C and D). The interacting residues correlate well with previous NMR chemical shift analysis and mutation studies regarding the charged and hydrophobic residues (Guler *et al*., 2012). On the left side of the groove, W517 of HelB fits into a well-defined pocket (named side-pocket thereafter) formed by RPA70 L45 and the aliphatic portions of N29, R31, R43 and S54 (Figure 1E and F). HelB residues D510, E516 and T519 were stabilized by hydrogen bonding or electrostatic interactions with the side chains of R31, N29 and R91. In addition, R43 forms a hydrogen bond with the main chain carbonyl group of W517, further stabilizing the folded-back conformation of the HelB peptide (Figure 1F). The overall binding mode of HelB to RPA70N is similar to that of p53 and DNA2 (Figure 1G). The amphipathic helix of HelB overlaps with one of the p53 peptides (Fig 1G, colored orange) and the DNA2 helix while the β turn coincides with part of the other p53 peptide (Fig 1G, colored yellow) and the β turn region of DNA2. All three peptides have a conserved hydrophobic residue that fits into the side pocket (Figure 1G). ITC titration results showed that mutation of W517 in HelB to alanine reduced the affinity between HelB peptide with RPA70N from around 4 µM to 16 µM (Figure 1H, Supplemental Figure S3), highlighting the contribution of side-pocket interactions to the overall binding strength.

**Fig. 1:**
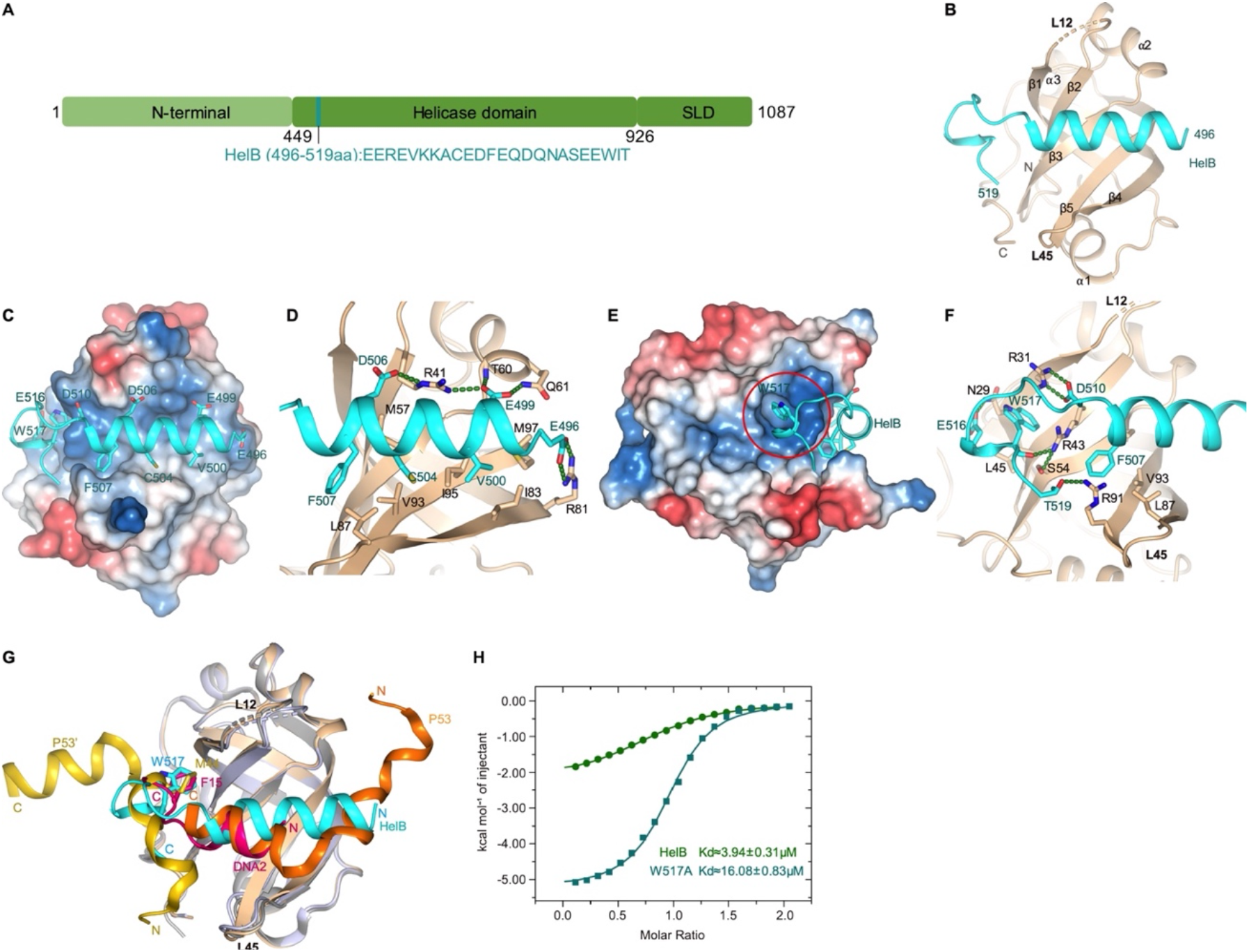
Structure of RPA70N-HelB complex. **A**. Linear domain diagram of HelB showing the position and sequence of RPA70N interacting motif. **B**. Ribbon representation of RPA70N-HelB crystal structure, HelB peptide is colored in cyan and RPA70N in beige. L12 denotes the loop between β1 and β2, L45 is the loop between β4 and β5. **C**. Same structure as in **B** showing the surface charge of RPA70N (red is negative, blue is positive), hydrophobic and negatively charged residues of HelB are displayed as sticks. **D**. Close-up view of the amphipathic HelB helix interacting with RPA70N groove, green dashed lines indicate hydrogen bonds or salt bridges. **E**. Side view of the RPA70N-HelB structure, the electrostatic surface of RPA70N is displayed and the side-pocket is highlighted, the representation is rotated 90° compared to **C**. **F**. Close-up view of the side-pocket residues coordinating the C-terminal part of HelB peptide, green dashed lines indicate hydrogen bonds or salt bridges. **G**. Superposition of RPA70N-HelB structure with RPA70N-p53 (PDB:2B3G) and RPA70N-DNA2 (PDB:5EAN) structures. HelB is colored in cyan, DNA2 in light-magenta, p53 in orange and yellow. The direction of HelB, p53 and DNA2 peptides in the RPA70N groove is the same. All three proteins have a hydrophobic residue inserted into the side-pocket of RPA70N. **H**. ITC titration results of WT HelB (496-519aa) or W517A mutant peptide with RPA70N.

### Structure of the RPA70N-ATRIP peptide complex

ATR is a member of the PIKK kinase family and ATR-ATRIP complex is a key regulator of DNA damage checkpoint, the complex is recruited to DNA damage sites by RPA coated ssDNA through ATRIP (Ball *et al*., 2005; Zou & Elledge, 2003). We crystallized RPA70N-ATRIP fusion protein in P212121 space group with one molecule in the asymmetric unit (Supplemental Table S2). ATRIP peptide binds to the RPA70N it fused to and the linker region is disordered. In the structure, ATRIP residues 53-68 form a 3-turn helix with 2 short flanking loops (Figure 2A and B, Supplemental Figure S2A). The hydrophobic side of the helix consisted of F55, L60, L63 and L66 packs against the broad hydrophobic patch of the RPA70N groove (Figure 2B). The N-terminus loop region of the ATRIP peptide is coordinated by R43 and R91 which form salt-bridges and hydrogen bonds with D54 and main chain carbonyl groups. At the C-terminus of the peptide, RPA70N R41 forms a hydrogen bond with the carbonyl group of L63 while ATRIP E62 forms a hydrogen bond with the main chain amide group of K88. The direction of the ATRIP peptide is inverted compared to HelB or DNA2 (Figures 2C, 1G), instead it is the same as seen in *K. lactis* Ddc2 (ATRIP)-Rfa1N complex (PDB: 5OMB)(Deshpande *et al*., 2017), both of which use a hydrophobic residue (F55 or I14) at the N-terminus to anchor the peptide at the groove (Figure 2C). Aiming to inhibit ATRIP-RPA70N interaction in cells and based on the structure of RPA70N-p53 complex, Frank et al. engineered a stapled helix peptide that binds to RPA70N and determined co-crystal structure of the synthetic helix with RPA70N (Frank *et al*, 2014). In their structure (PDB:4NB3), the peptide is in a reversed orientation compared to our structure or the Ddc2-Rfa1N structure and employs a 3,4-dichloro-substituted phenylalanine (ZCL) to bind the hydrophobic pocket where F55 in ATRIP binds (Figure 2D). Mutation of F55 to alanine greatly reduced the affinity of ATRIP towards RPA70N (Figure 2E, Supplemental Figure S4), indicating the hydrophobic interactions mediated by F55 is critical for maintaining ATRIP-RPA70N association.

**Fig. 2:**
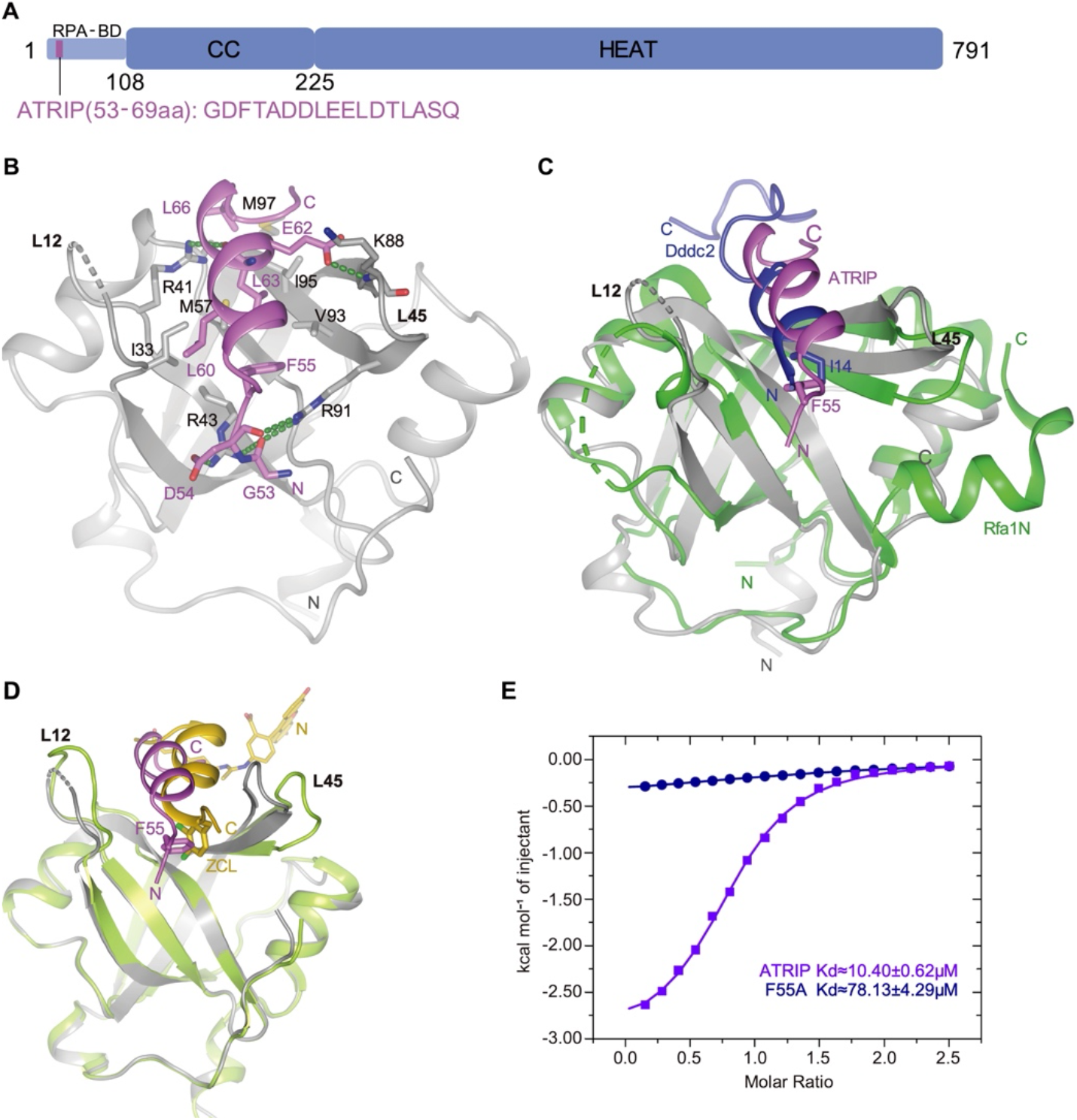
Structure of RPA70N-ATRIP complex. **A**. Linear domain diagram of ATRIP showing the position and sequence of RPA70N interacting motif. **B**. Ribbon representation of RPA70N-ATRIP crystal structure, ATRIP peptide is colored in violet and RPA70N in light grey. Important interacting residues are shown as sticks, green dashed lines indicate hydrogen bonds or salt bridges. **C**. Alignment of RPA70N-ATRIP structure with Ddc2-Rfa1N structure (PDB: 5OMB) showing that ATRIP and Ddc2 bind to RPA70N in the same direction. RPA70N-ATRIP are colored as in **B**, Ddc2 is colored in green and Rfa1N is colored in blue. **D**. Superposition of RPA70N-ATRIP structure with RPA70N-stapled peptide complex (PDB:4NB3), for 4NB3 RPA70N is colored in light-green and the stapled peptide is colored in yellow. ZCL is 3,4-dichloro-substituted phenylalanine. The direction of the stapled peptide is reversed compared to ATRIP. **E**. ITC titration results of WT ATRIP peptide (53-69aa) or F55A mutant peptide with RPA70N.

### Structures of the RPA70N-BLM peptide complexes

BLM helicase is a multifunctional RecQ family helicase, it is involved in DNA-end resection, restart of stalled replication forks, dissolving Holliday junctions, and processing of ultra-fine DNA bridges (Bythell-Douglas & Deans, 2021; Chu & Hickson, 2009; Croteau *et al*, 2014; Kitano, 2014; Shorrocks *et al*., 2021). It has two RPA70N binding motifs in the N-terminal disordered region, namely residues 146-165 (BLMp1) and residues 550-570 (BLMp2)(Doherty *et al*., 2005; Shorrocks *et al*., 2021). We fused BLMp1 and BLMp2 separately to RPA70N and determined their structures (Supplemental Table S2 and S3).

In the structure of RPA70N-BLMp2, BLM residues 550-564 are ordered and coordinated by two RPA70N molecules (Figure 3A and B, Supplemental Figure S2B). The C-terminal part of the kinked peptide fits onto the RPA70N groove, with F556, I558 and F561 making contacts with the hydrophobic patch of RPA70N while RPA70N residues R41, K88, R91 and R43 form salt bridges or hydrogen bonds with D560, D552, D562 and the main chain carbonyl group of F561 (Figure 3C). Interestingly, the N-terminal half of BLMp2 latches onto the α1 region of a nearby RPA70N (Figure 3D). K16’ from RPA70N forms several ionic interactions with D554, D557 and D559 to neutralize the negative charges. Q15’ also contributes to the interaction by forming two hydrogen bonds with BLM D557. Near the tip of the BLMp2 peptide, Y551 fits onto a small hydrophobic surface formed by A9’, A12’, I13’ and I21’, its main chain amide group also forms a hydrogen bond with the carbonyl group of E7’ (Figure 3D). Overall, it appears BLMp2 promoted Loop 12 of one RPA70N to interaction with α1’ of the other RPA70N and each RPA70N provides some of the binding surface for BLMp2 (Figure 3D, Supplemental Figure S5A). With ITC, we found that BLMp2 binds to RPA70N with a relatively weak *K*_*D*_ around 18 µM (Figure 3E, Supplemental Figure S6A).

**Fig. 3:**
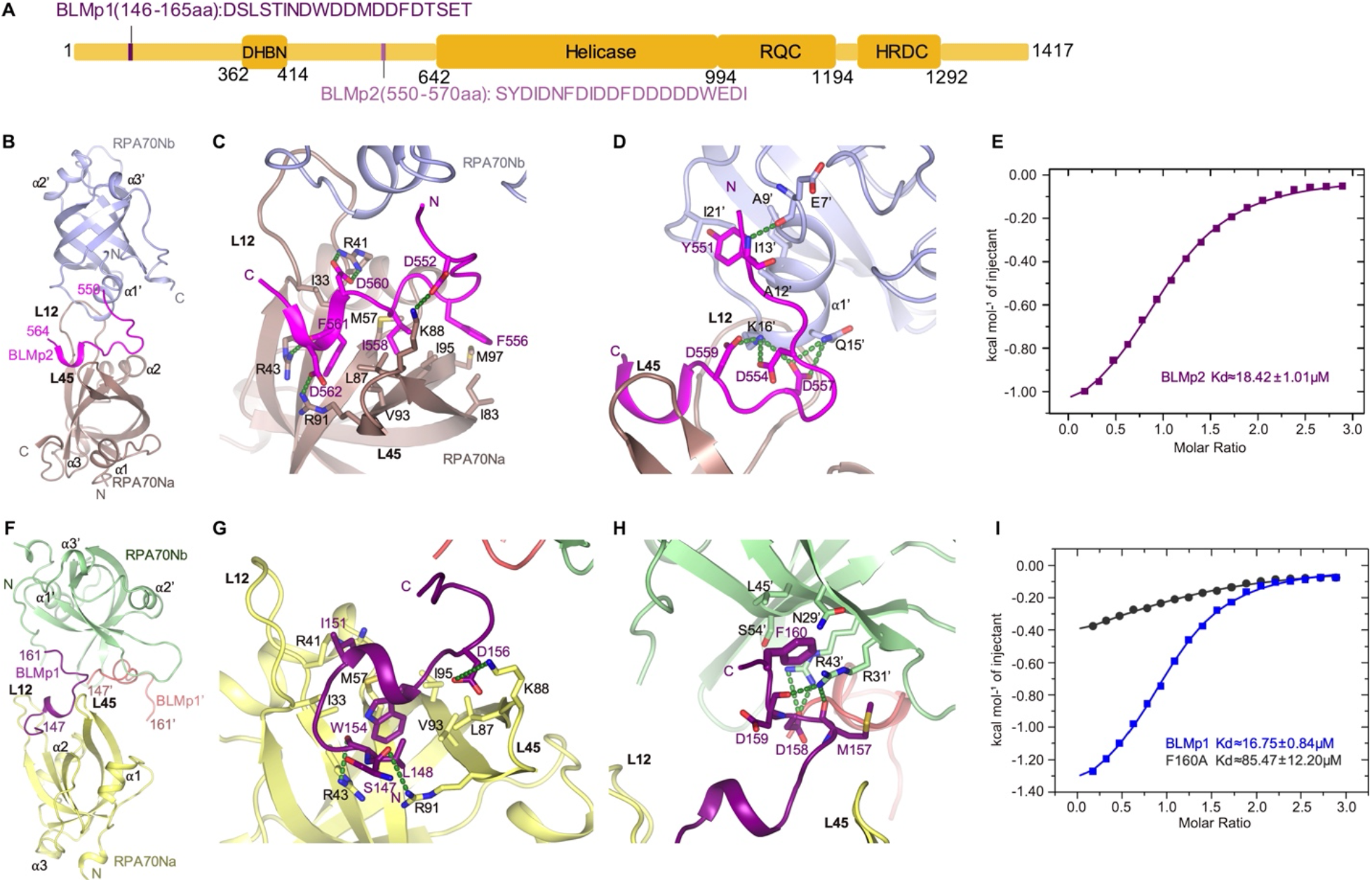
Structures of two RPA70N-BLM complexes. **A**. Linear domain diagram of BLM showing the position and sequence of RPA70N interacting motifs. **B**. Ribbon representation of RPA70N-BLMp2 crystal structure, the BLMp2 peptide is coordinated by two RPA70N molecules. BLMp2 is colored in magenta and the two RPA70N molecules are colored in chocolate and light-blue, respectively. The other BLMp2 peptide in RPA70Nb is colored pink. **C**. Close-up view of BLMp2 interacting with the RPA70Na groove, interacting residues are shown as sticks, green dashed lines indicate hydrogen bonds or salt bridges. **D**. Close-up view of BLMp2 interacting with RPA70Nb α1, interacting residues are shown as sticks, green dashed lines indicate hydrogen bonds or salt bridges. **E**. ITC titration result of WT BLMp2 peptide (550-570aa) with RPA70N. **F**. Ribbon representation of RPA70N-BLMp1 crystal structure, the BLMp1 peptide is coordinated by two RPA70N molecules. BLMp1 is colored in purple and the two RPA70N molecules are colored in yellow and light-green, respectively. **G**. Close-up view of BLMp1 interacting with the RPA70Na groove, interacting residues are shown as sticks, green dashed lines indicate hydrogen bonds or salt bridges. **H**. Close-up view of BLMp1 interacting with RPA70Nb side-pocket, interacting residues are shown as sticks, green dashed lines indicate hydrogen bonds or salt bridges. **I**. ITC titration results of WT BLMp1 peptide (146-165aa) or F160A mutant peptide with RPA70N.

In the structure of RPA70N-BLMp1, BLMp1 also adopts a kinked conformation and is coordinated by two RPA70N molecules (Figure 3F, Supplemental Figure S2C). However, one major difference is that BLMp1 binds to RPA70N in a reversed direction compared to BLMp2, HelB or DNA2 (Figures 3F-G and 1G). The N-terminal part of BLMp1 forms a one-turn helix followed by a γ turn (Figure 3G). BLM W154 inserts into a hydrophobic pocket formed by RPA70N V93, I95, M57, I33 and the aliphatic part of R43 and stacks with M57. BLM L148 stacks on top of W154 and BLM I151 packs onto the side chains of RPA70N I33, M57 and R41 (Figure 3G). At the middle of the BLMp1 peptide, D156 is coordinated by K88 and the peptide forms another β turn. The C-terminal part of BLMp1 adopts an extended conformation, with F160 anchored in the side-pocket of a nearby RPA70N’ (Figure 3H). The RPA70N’ residues R43’, R31’ also interact with D158 side chain and main chain oxygen atoms of D159 and M157. The *K*_*D*_ of the BLMp1-RPA70N complex, determined by ITC, is around 16.7 µM, similar to BLMp2 (Figure 3I, Supplemental Figure S6B). Mutation of F160 to alanine greatly reduced the affinity between BLMp1 and RPA70N, resulting in a *K*_*D*_ around 85 µM (Figure 3I, Supplemental Figure S6C). The two RPA70N molecules are connected by the BLMp1 peptide but not making other contacts (Figure 3F, Supplemental Figure S5B). We could also present the structure as one RPA70N bound to two peptides as reported for the p53 peptide (Supplemental Figure S5C). As mentioned earlier, the direction of the BLMp1 peptide in the groove is reversed compared to the p53 peptide, the direction of the BLMp1’ peptide bound to the side pocket is the same as p53’ (Supplemental Figure S5C).

### Structure of the RPA70N-RMI1 peptide complex

RMI1 is another RPA partner, it is a subunit in the BTR complex and mainly mediates protein-protein interaction (Shorrocks *et al*., 2021; Wang *et al*, 2010; Xu *et al*, 2008a; Xue *et al*., 2013).The RPA70N interaction motif is located between its two OB folds (Figure 4A). In the RPA70N-RMI1 complex structure, RMI1 residues 243 and 259 form two short helixes with a β turn in the middle (Figure 4B, Supplemental Figure S2D). The overall arrangement is similar to the complex of RPA70N-BLMp1 with the N-terminal helix in the groove and the C-terminal helix binding to a neighboring RPA70N’ (Figures 4B, 3F). RMI1 L247 of the N-terminal helix fits into a hydrophobic pocket at the bottom of the RPA70N groove (Figure 4C). L248 and L251 interact with the hydrophobic side chains of RPA70N I33, M57 and the aliphatic part of R41. RMI1 D244 and E246 are stabilized by electrostatic interactions with R31, R43 and R91. The folded-back C-terminal helix also interacts with K88 by forming several hydrogen bonds (Figure 4C). RMI1 N254 inserts into the side-pocket of a neighboring RPA70N’ and forms a few hydrogen bonds with N29’ and R31’ side chains (Figure 4D). D252 also interacts with S54’ and R43’, further strengthening the interaction (Figure 4D). Analogous to RPA70N-BLMp1, the two RPA70N molecules coordinating RMI1 peptide aren’t making any contacts (Supplemental Figure S5D). Superposition of the two structures showed that F160 and N254 point to the same direction but not at the exact same location, indicating the second RPA70N molecule could adjust to different peptide sequences for binding (Figure 4E). ITC titration showed that RMI1 peptide binds to RPA70N with a *K*_*D*_ around 14.5 µM, mutation of N254 reduced the affinity between RMI1 and RPA70N to around 25.6 µM (Figure 4F, Supplemental Figure S7), which suggests the side chain of N254 probably doesn’t make as much contribution to the binding affinity as hydrophobic residues at this position, other interactions mediated by D252 or E256 might help to stabilize RMI1-RPA70N contacts (Figure 4C and D).

**Fig. 4:**
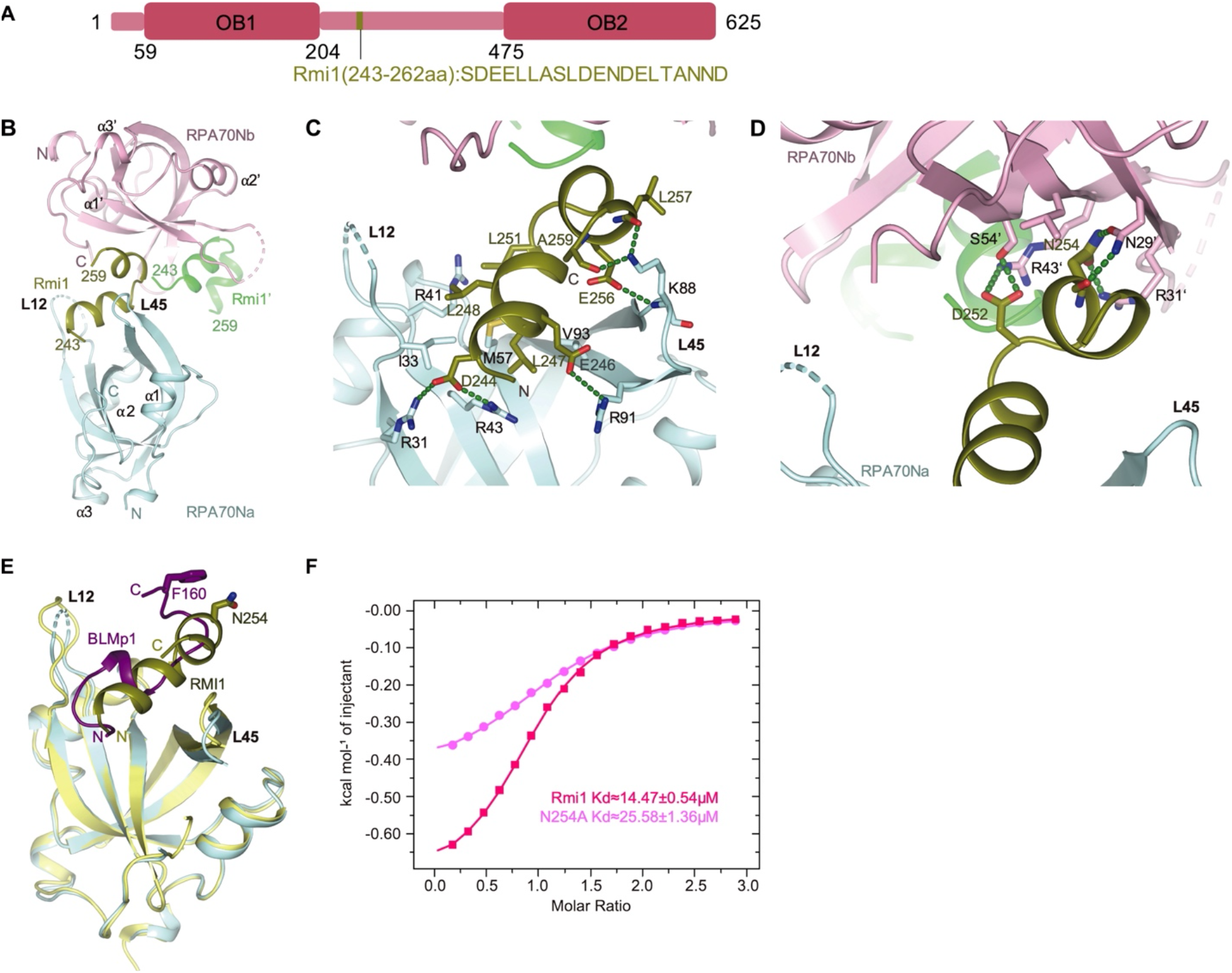
Structure of RPA70N-RMI1 complex. **A**. Linear domain diagram of RMI1 showing the position and sequence of RPA70N interacting motif. **B**. Ribbon representation of RPA70N-RMI1 crystal structure, the RMI1 peptide is coordinated by two RPA70N molecules. RMI1 is colored in olive and the two RPA70N molecules are colored in pale-cyan and light-pink respectively. The other RMI1 peptide in RPA70Nb is colored green. **C**. Close-up view of RMI1 interacting with the RPA70Na groove, interacting residues are shown as sticks, green dashed lines indicate hydrogen bonds or salt bridges. **D**. Close-up view of RMI1 interacting with RPA70Nb side-pocket, interacting residues are shown as sticks, green dashed lines indicate hydrogen bonds or salt bridges. **E**. Superposition of RPA70N-RMI1 structure with RPA70N-BLMp1 complex, RMI1 and BLMp1 interact with RPA70N in a similar manner. **I**. ITC titration results of WT RMI1 peptide (243-262aa) or N254A mutant peptide with RPA70N.

### Structure of the RPA70N-WRN peptide complex

WRN nuclease-helicase belongs to the RecQ family of DNA helicases and play important roles in DNA repair and maintenance of genome integrity (Chu & Hickson, 2009; Croteau *et al*., 2014; Kitano, 2014; Mukherjee *et al*, 2018). WRN has two tandem RPA binding motifs with the same sequence localized between its nuclease and helicase domains (Doherty *et al*., 2005; Shen *et al*., 2003; Yeom *et al*., 2019) (Figure 5A). We fused one WRN motif to RPA70N and the fusion construct crystalized in space group P212121 with two molecules in the asymmetry unit (Supplemental Table S3). In the structure, WRN 435 to 451 formed a continuous helix and inserts into the amphipathic groove of a symmetry related RPA70N (Figure 5B). Most of the WRN helix is coordinated by residues in the groove, specifically E439-R31’-D443-R43’-E442-R91’-E445 form a series of electrostatic interactions, and M446, L449 contact the hydrophobic patch formed by L87’, V97’, I33’, M57’ and I95’ (Figure 5C). The N-terminal part of the each WRN peptide helix interacts with the RPA70N it fused to. Y436’ fits into the side-pocket and forms two hydrogen bonds with R31’ (Figure 5D). Compared to the RPA70N-p53 structure, the direction of the WRN peptide in the groove is reversed and the position of the residues (Y436 or M44) to interact with the side pocket is quite different (Figure 5E). The direction of the other WRN’ peptide bound to the side pocket is the same as p53’ (Figure 5E). ITC titration of WRN peptide with RPA70N yielded a *K*_*D*_ around 11.6 µM while M466A mutation increased *K*_*D*_ value to around 37.4 µM (Figure 5F and G, Supplemental Figure S8).

**Fig. 5:**
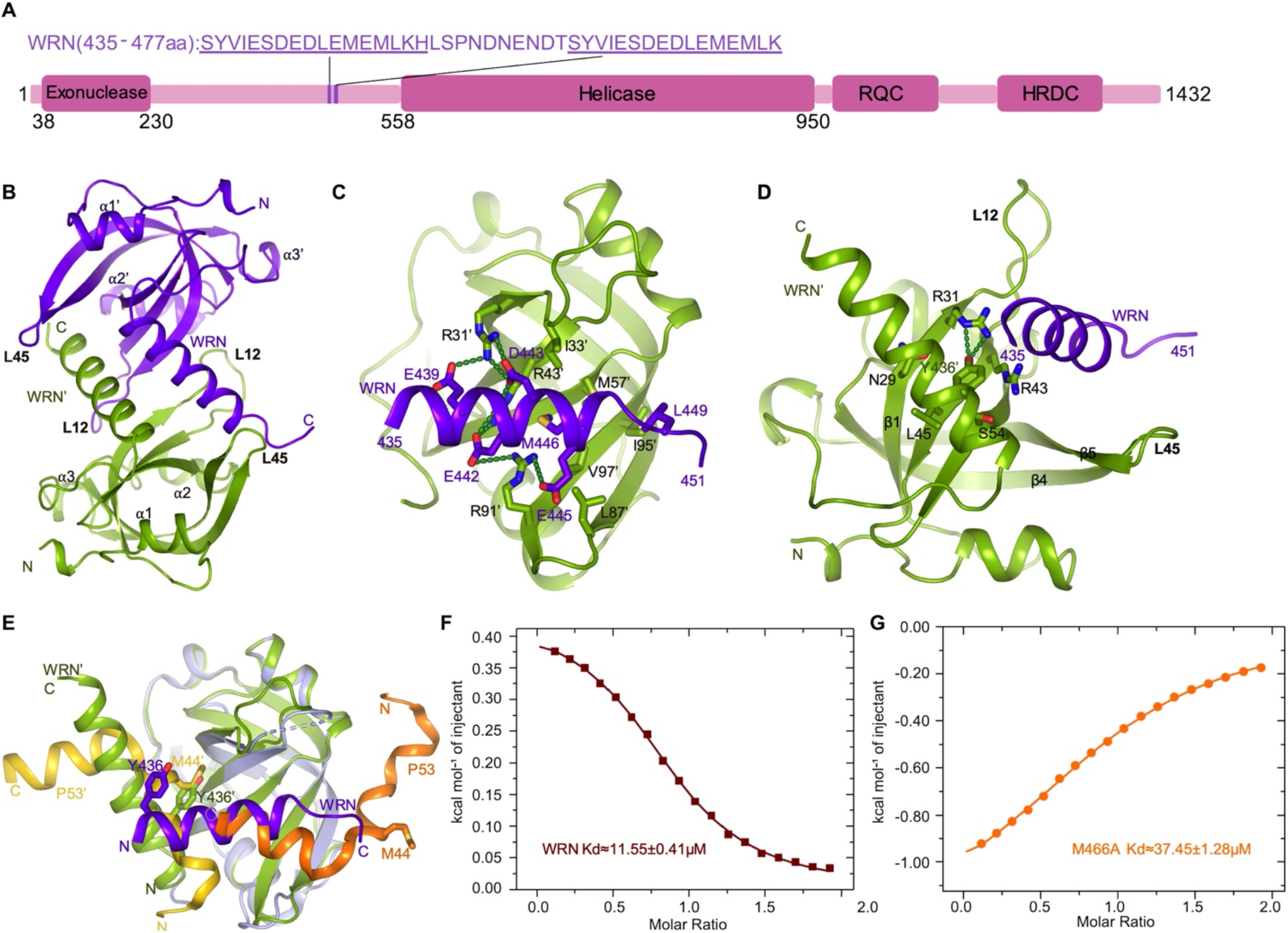
Structure of RPA70N-WRN complex. **A**. Linear domain diagram of WRN showing the position and sequence of RPA70N interacting motifs. **B**. Ribbon representation of RPA70N-WRN crystal structure, the fused WRN peptide forms an α helix and inserts into the groove of the symmetry-related RPA70N molecule. The two RPA70N molecules and linked WRN peptides are colored in purple-blue and light-green, respectively. **C**. Close-up view of WRN interacting with the RPA70N’ groove, interacting residues are shown as sticks, green dashed lines indicate hydrogen bonds or salt bridges. **D**. Close-up view of WRN’ interacting with RPA70N’ side-pocket, interacting residues are shown as sticks, green dashed lines indicate hydrogen bonds or salt bridges. **E**. Superposition of RPA70N-WRN structure with RPA70N-p53 complex (PDB:2b3G), WRN’ (Y436’) binds to the side-pocket similarly to p53’ (M44’), while Y436 in WRN and M44 in p53 are in different locations due to the inverted orientation of WRN and p53 in RPA70N groove. **F**. ITC titration result of WT WRN peptide (435-451aa) with RPA70N, the titration appears to be endothermic. **G**. ITC titration result of M466A mutant WRN peptide (435-451aa) with RPA70N.

## Discussion

It is well established that RPA plays important roles in DNA replication, recombination and repair (Caldwell & Spies, 2020; Chen & Wold, 2014; Fanning *et al*., 2006; Iftode *et al*., 1999; Marechal & Zou, 2015; Wold, 1997; Zou *et al*., 2006). Many of the RPA-protein interactions are mediated by the flexibly tethered RPA70N domain. However, RPA70N-partner interactions are often weak and highly dynamic (Caldwell & Spies, 2020; Fanning *et al*., 2006). As a result, high resolution structures of RPA70N bound to partner peptides are rare compared to the many protein RPA70N interacts with. To overcome this problem, inspired by the fusion approach first employed by Bochkareva et al. to solve the RPA70N-p53 complex structure (Bochkareva *et al*., 2005), we systematically screened RPA70N-partner fusion constructs for crystallization and determined 6 complex structures of proteins involved in DNA damage response (Supplemental Table S1). These structures confirmed previous findings that RPA70N binds to a partner sequence through two interfaces, one is the basic and hydrophobic groove, the other is the side-pocket which is also basic and hydrophobic. The side-pocket is not always used as seen in the cases of ATRIP, BLMp2 and Primpol. In theory, the empty side-pocket could be the binding site of another peptide which is able to binds there. This second peptide could be a not-yet identified sequence in the protein RPA70N bound to or from another molecule. More importantly, we found that RPA70N could coordinate peptide binding to its two interfaces through diverse means, e.g., inverted direction, rotation/tilt of the bound helix, kinked conformation, dimerization, etc. The versatile ways of interaction are presumably customized to the different protein sequences RPA encounters. One could imagine that RPA70N must be able to recruit different partners under different scenarios.

Of particular interest is that many of the partner peptides appear to be able to connect two RPA70N domains (Supplemental Figures S5A, B and D). If we expand the dimer observed, we could get a string of RPA70N connected by BLMp1, BLMp2 or RMI1. Intriguingly, some of these partner proteins themselves (BTR complex, WRN, ATRIP, p53) are often dimers or oligomers (Cho *et al*, 1994; Compton *et al*, 2008; Deshpande *et al*., 2017; Hodson *et al*, 2022). So there are indeed multiple copies of partner peptide in close range. One RPA usually covers around 30nt ssDNA (Kim *et al*., 1994), for medium to long ssDNA there are multiple copies of RPA bound. The bridge forming nature of the partner peptides in combination with many RPAs on ssDNA could greatly enhance the efficiency of partner recruitment when needed, for example, in DNA damage response (Figure 6).

**Fig. 6:**
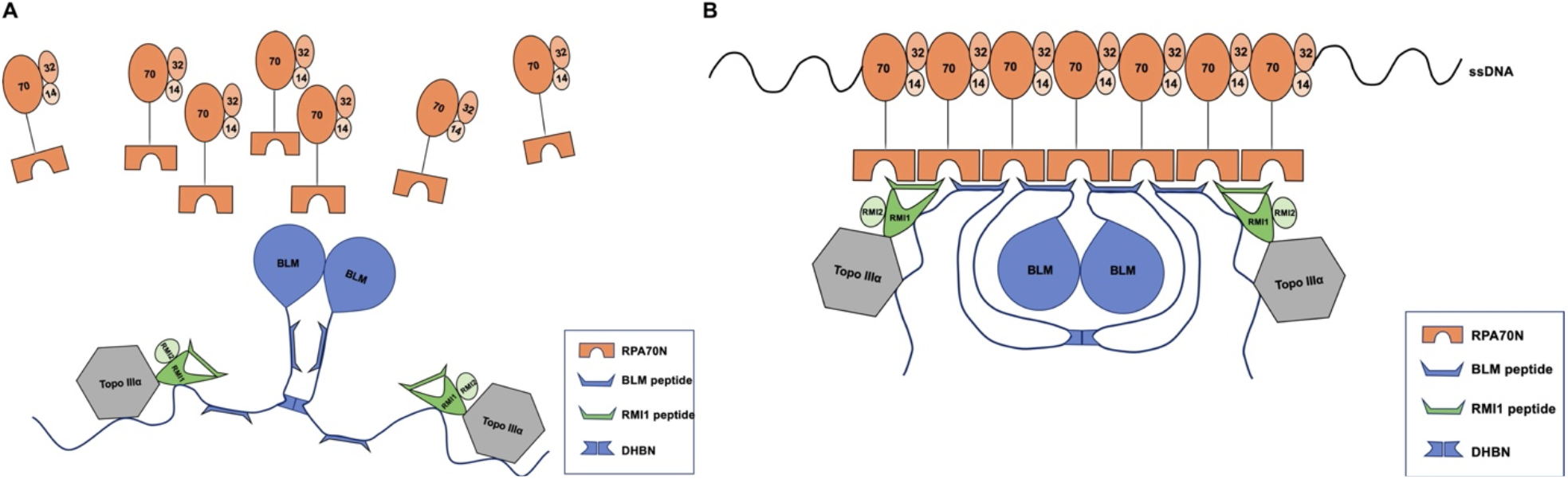
A model of RPA-ssDNA mediated BTR complex recruitment. **A**. Without ssDNA, RPA molecules are not associated with the BTR complex due to relative weak affinity between RPA70N and the interacting peptide sequences in the BTR complex. **B**. Multiple RPA molecules nucleate on ssDNA, the increased avidity of RPA70N promotes BTR peptides to bind, bound peptides in turn promotes RPA70N association.

The multivalent way of interaction could also serve as an intrinsic layer of regulation in addition to signal transduction pathways like protein phosphorylation. Under normal conditions RPA molecules are not clustered on ssDNA and have relative weak affinity for many DNA damage response proteins as shown by the dissociation constant values measured in this study and previous studies (Hegnauer *et al*, 2012; Lee *et al*, 2018; Souza-Fagundes *et al*, 2012; Yeom *et al*., 2019). With DNA damage, RPAs nucleate on exposed ssDNA quickly due to their sub nanomolar affinity for ssDNA and recruit corresponding proteins: with sufficient copies of RPA bound to ssDNA, the weak affinity of monomeric RPA70N toward target protein is now overwhelmed by multiple interaction interfaces (Figure 6). Shorrocks and coworkers reported that within 5 min RPA and BLM proteins accumulated rapidly to laser lines but at relatively low levels, RPA microfoci appeared in ∼50% of irradiated cells after 15 min as a result of DNA end-resection in S and G2 cells, shortly after at 20 min BLM microfoci appeared and co-localized with RPA while a BLM mutant lacking both BLMp1 and BLMp2 failed to form microfoci (Shorrocks *et al*., 2021). Studies carried out by Doherty et al. and Lee et al. showed that RPA stimulates WRN helicase activity in a concentration dependent manner and the helicase activity of WRN requires the binding of multiple RPAs (Doherty *et al*., 2005; Lee *et al*., 2018). In addition, a recent study revealed that all ssDNA-dependent activities of HelB are greatly stimulated by RPA-ssDNA filaments (Hormeno *et al*., 2022).

Since the physical association of RPA70N with DNA damage response proteins are critical for maintaining genome integrity, it is also subjected to negative regulation. The N-terminus of RPA32 could be hyperphosphorylated by DNA-PK, ATM, and ATR, which in turn binds to RPA70N and inhibits RPA70N mediated protein recruitment (Binz *et al*, 2003; Binz & Wold, 2008; Lee *et al*, 2020). Hyperphosphorylated RPA32 disrupted RPA-p53, RPA-MRN interactions (Abramova *et al*., 1997; Bochkareva *et al*., 2005; Oakley *et al*., 2009). In vitro analysis showed that controlling access to RPA70N through phosphorylation of RPA32 by ATR after DNA damage may be a mechanism for regulating cell cycle progression without impairing DNA repair (Lindsey-Boltz *et al*, 2012). A recent study carried out by Soniat et al. demonstrated that RPA70N binding to phosphorylated RPA32 suppressed BLM helicase interaction and inhibited DNA end resection (Soniat *et al*, 2019). Similar mechanisms are highly likely to exist for other proteins that RPA70N binds. It would be interesting to find out the detail mechanism of how phosphorylated RPA32 compete and abrogate RPA70N interactions with a variety of DNA damage response proteins in the future.

In short summary, the structural snapshots and biochemical analysis we present here shed light on the diverse modes of RPA70N interacting with DNA damage response proteins and these interactions could serve to increase the avidity of RPA70N binding.

## Materials and Methods

### Cloning, protein expression and purification

The DNA sequences of human RPA70N (residue 1-120), HelB (residue 496-519), ATRIP (residue 53-69), BLMp1 (residue 146-165), BLMp2 (residue 550-570), RMI1 (residue 243-262) and WRN (residue 435-451) were cloned into a modified pRSFDuet-1 vector (Novagen) which fuses an N-terminal 6-His-sumo tag to the target gene using ClonExpress II One Step Cloning Kit (Vazyme). RPA70N-peptide fusion constructs were cloned into the same expression vector. RPA70N, RPA70N-peptide fusion proteins and all peptides were expressed and purified with similar steps. The recombinant plasmids were transformed into *E. coli* BL21(DE3) cells (Novagen), which were grown in LB medium at 37°C until the OD 600 reached 0.6–0.8. Overexpression of proteins were induced by addition of 0.5 mM isopropyl β-D-thiogalactopyranoside (IPTG), followed by incubation at 20°C for 14 h. Cells were harvested by centrifugation, resuspended in lysis buffer (20 mM Tris-HCl, 200 mM NaCl, 20 mM imdazole, 10% glycerol, 0.3 mM TCEP, pH 8.0), and lysed by a high-pressure homogenizer at 4°C. The cell lysate was centrifuged at 12000 rpm for 40 min to obtain soluble extract. After nickel affinity pull-down, 6-His-sumo tag was cleaved off by Ulp1 protease and removed by a second nickel column. Flow-through was then passed through a Source 15Q column (Cytiva) and eluted with a gradient of 0-1 M NaCl in a buffer of 20 mM Tris-HCl, pH 8.0, 10% glycerol, 0.3 mM TCEP. Fractions containing target proteins were pooled and concentrated, then further purified on a Superdex 75 increase gel filtration column (Cytiva) in a buffer containing 20 mM Tris-HCl, pH 8.0, 150 mM NaCl, 0.3 mM TCEP. The purified RPA70N-peptide fusion proteins were concentrated to around 20-25 mg/ml for crystallization. RPA70N and all peptides were concentrated to suitable concentrations for ITC titrations.

### Isothermal titration calorimetry (ITC)

All ITC titrations were carried out using a MicroCal PEAQ-ITC instrument (Malvern) at 25°C with different peptides in the syringe and RPA70N in the cell. RPA70N and peptide samples were dialyzed against a working buffer consisting of 20 mM HEPES, 100 mM NaCl, 1 mM DTT, pH 7.5. Partner peptides at 1.0 mM were titrated into RPA70N at 0.1 mM in the well. Each titration was carried out with 19 injections spaced 150s apart, with 0.4 μl used for the first and 2.4 μl used for the rest. The acquired calorimetric titration data were analyzed with Origin 7.0 software using the ‘One Set of Binding Sites’ fitting model.

### Crystallization

For all the RPA70N-peptide fusion proteins, crystallization screenings were performed using 96-well plates in a sitting drop mode at 4°C. The RPA70N-HelB fusion protein crystallized in 20% (w/v) PEG 3350, 200 mM calcium chloride. The RPA70N-BLMp1 fusion protein crystallized in 100 mM sodium citrate pH 5.6, 2000 mM ammonium sulfate, 200 mM potassium/sodium tartrate. RPA70N-BLMp2 fusion protein crystallized in 100 mM sodium acetate pH 4.6, 8% (w/v) PEG 4000. The RPA70N-ATRIP fusion protein crystallized in 100 mM Tris-HCl pH 8.5, 2400 mM ammonium sulfate. The RPA70-WRN fusion protein crystallized in 20% (w/v) PEG 3350, 200 mM ammonium sulfate. The RPA70N-RMI1 fusion protein crystallized in 100 mM sodium acetate pH 4.6, 30% (w/v) PEG 2000 MME, 200 mM ammonium sulfate. Crystals were cryo-protected in their respective well solutions supplemented with 20% ethylene glycol and flash-frozen in liquid nitrogen.

### Structure determination and refinement

Diffraction data were collected at Beamline stations BL17U1, BL18U1 and BL19U1 at Shanghai Synchrotron Radiation Facility (SSRF, Shanghai, China). The data were integrated and scaled using XDS, the CCP4 program Pointless and Aimless (Evans & Murshudov, 2013; Kabsch, 2010; Winn *et al*, 2011). The structures of RPA70N-peptide fusion constructs were determined by molecular replacement using the RPA70N structure from PDB 5EAY as an initial searching model with Phaser (McCoy *et al*, 2007). The structural model was built using Coot (Emsley & Cowtan, 2004) and refined using PHENIX (Liebschner *et al*, 2019). Figures were generated using PyMOL (The PyMOL Molecular Graphics System, Version 2.0 Schrödinger, LLC). The statistics of the data collection and refinement are shown in Supplemental Table S2 and S3.

## Data Availability

Atomic coordinates and structure factors for the reported crystal structures have been deposited with the Protein Data Bank under accession number: 7XUT, 7XUV, 7XUW, 7XV0, 7XV1 and 7XV4.

## Acknowledgements

We thank the staff at SSRF BL19U1, BL17U1 and BL18U1 for X-ray diffraction data collection. We thank Yuyuan Zheng, Panyu Fei and all the members of Zhou Lab for their kind help. This work was supported by National Natural Science Foundation of China (31971125 to C.Z.).

## Author contributions

Y.W., N.Z. and W.F. cloned, expressed and purified proteins. Y.W. and W.F. performed ITC titrations. Y.W., N.Z. and W.F. carried out crystallization experiments and collected the crystallographic data. C.Z. carried out structural determination and analysis. C.Z. and Y.W. wrote the manuscript.

## Supplementary Information for

**Fig. S1:**
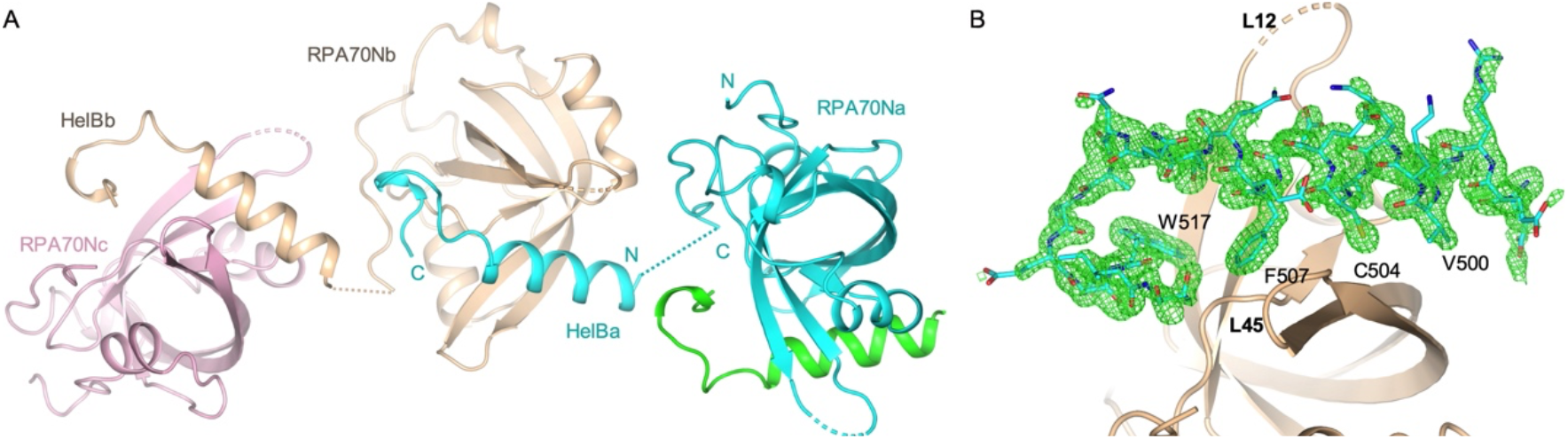
Analysis of RPA70N-HelB complex structure. **A**. Crystal packing analysis indicates that HelB peptide from one RPA70N-HelB fusion molecule binds to the RPA70N portion of a neighboring molecule in the crystal. **B**. *mFo-DFc* electron density map (green mesh) of HelB peptide contoured at 2.0 σ.

**Fig. S2:**
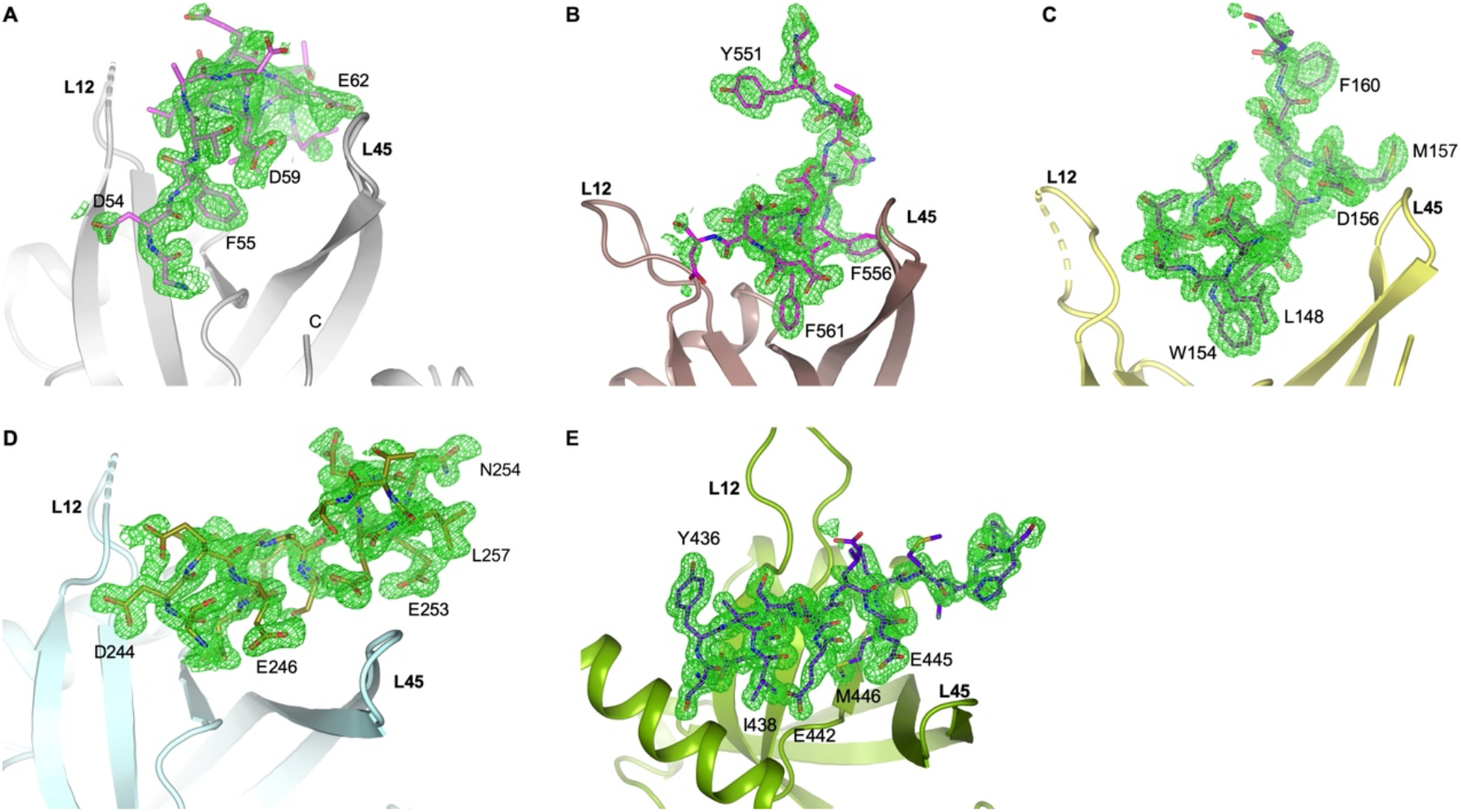
Electron density omit map of RPA70N-peptide complexes. **A**. *mFo-DFc* electron density map (green mesh) of ATRIP peptide contoured at 2.0 σ. **B**. *mFo-DFc* electron density map (green mesh) of BLMp2 peptide contoured at 2.0 σ. **C**. *mFo-DFc* electron density map (green mesh) of BLMp1 peptide contoured at 2.0 σ. **D**. *mFo-DFc* electron density map (green mesh) of RMI1 peptide contoured at 2.0 σ. **E**. *mFo-DFc* electron density map (green mesh) of WRN peptide contoured at 2.0 σ.

**Fig. S3:**
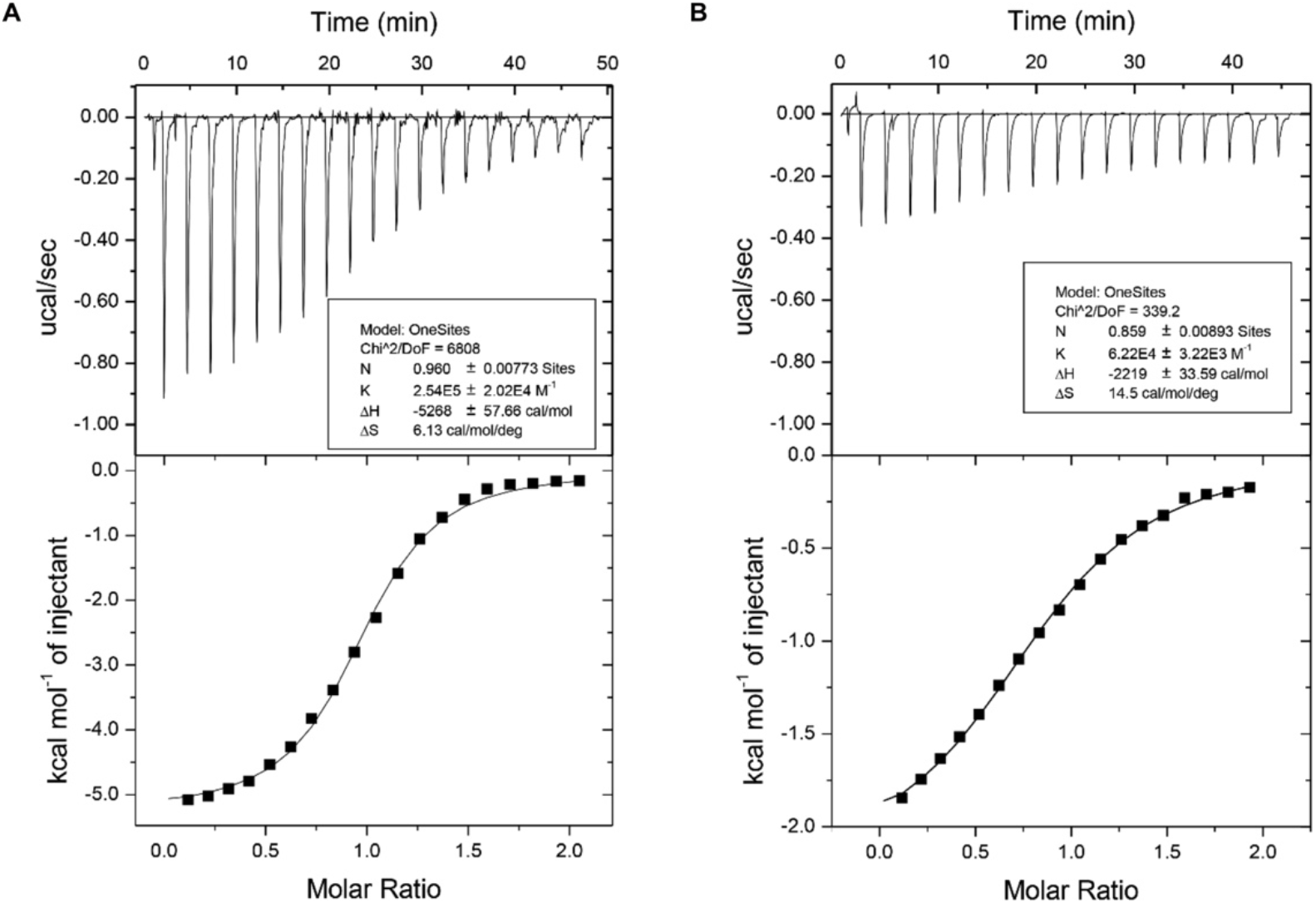
ITC titration profiles of (A) WT HelB (496-519aa) peptide or (B) W517A mutant peptide with RPA70N.

**Fig. S4:**
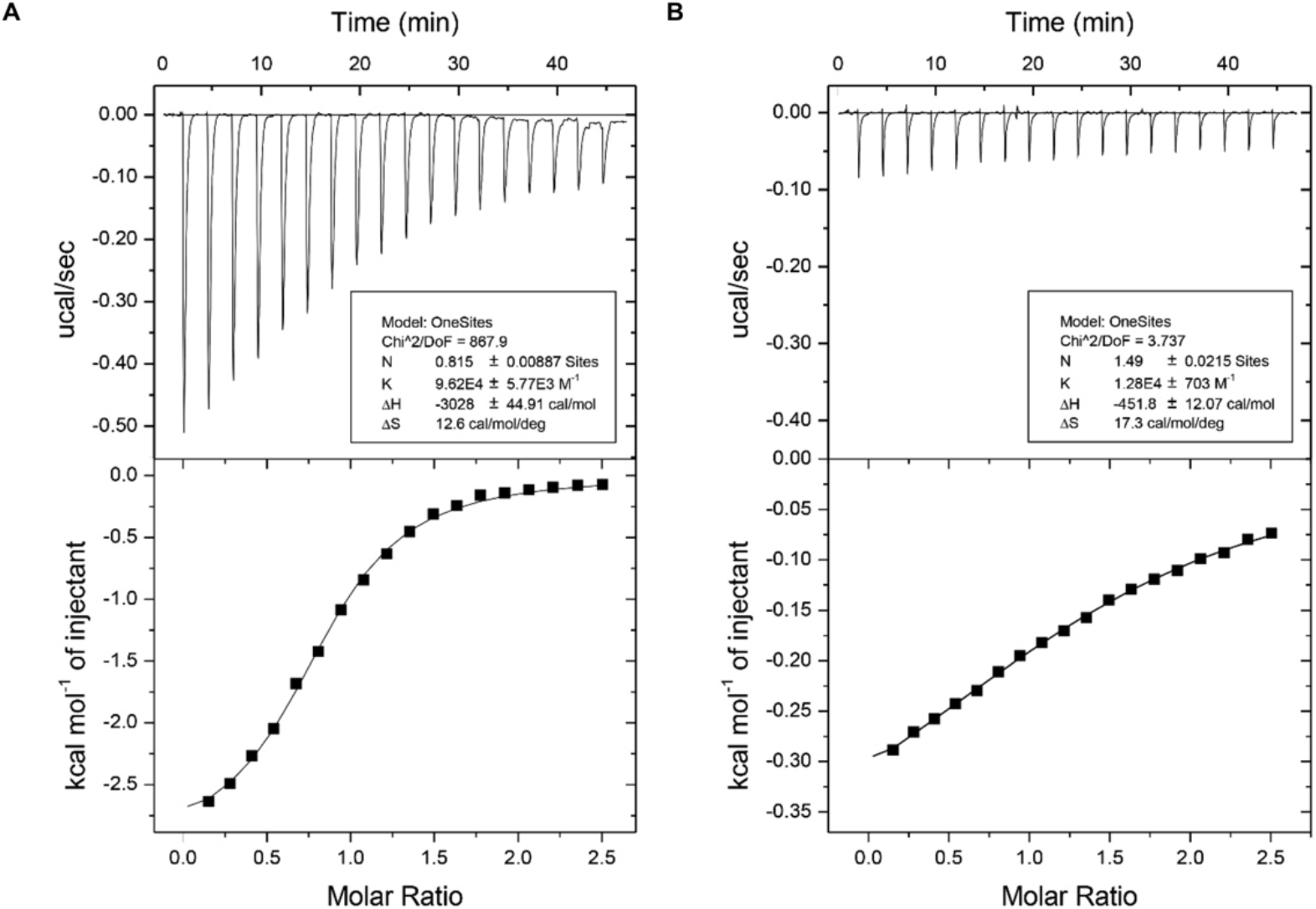
ITC titration profiles of (A) WT ATRIP peptide (53-69aa) or (B) F55A mutant peptide with RPA70N.

**Fig. S5:**
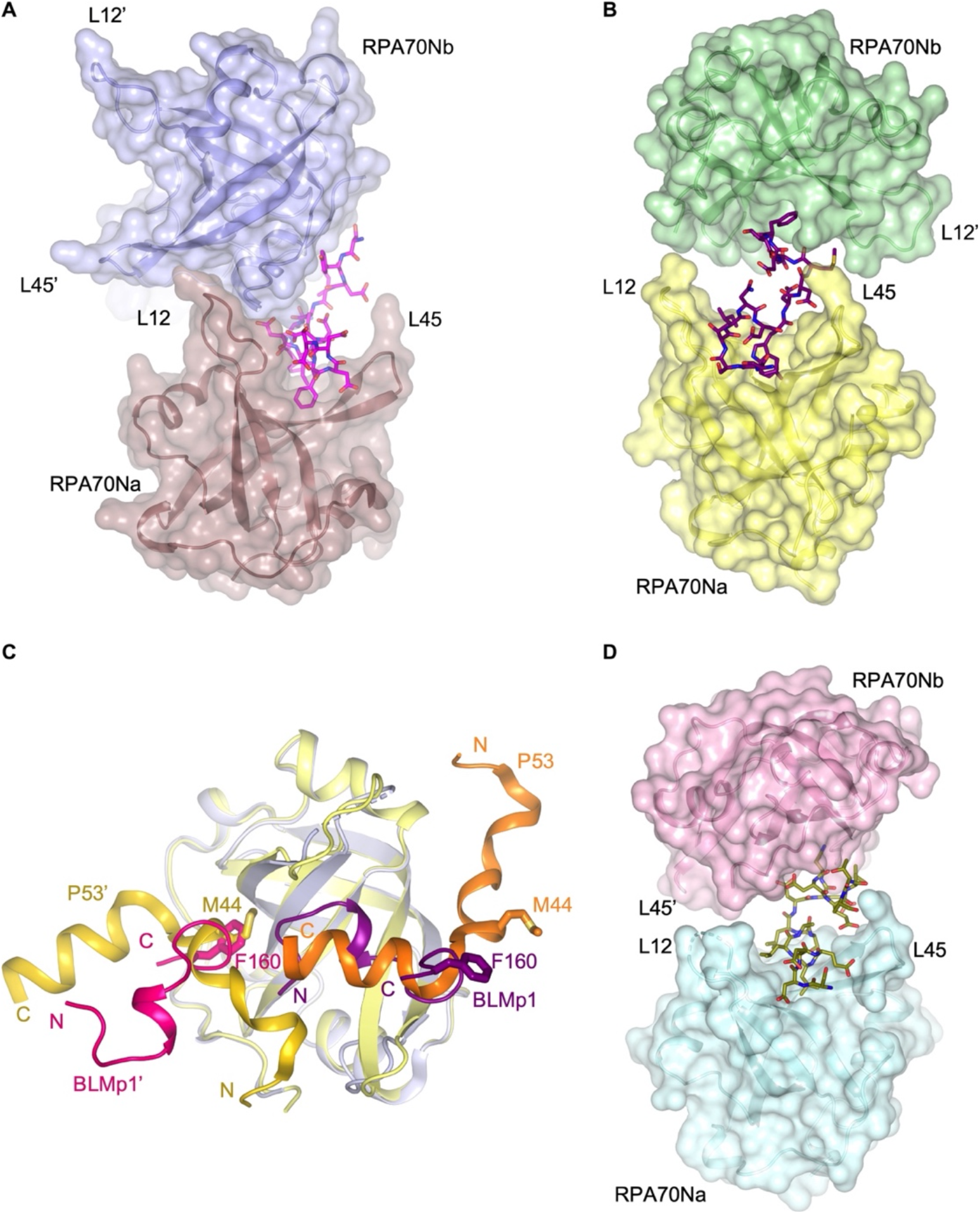
Analysis of BLM, RMI1 peptide-RPA70N complex structures. **A**. Surface representation of the two RPA70N molecules interacting with BLMp2 peptide which is displayed as magenta-colored sticks. **B**. Surface representation of the two RPA70N molecules interacting with BLMp1 peptide which is displayed as purple-colored sticks. **C**. Superposition of RPA70N-BLMp1 structure with RPA70N-p53 complex (PDB:2B3G), showing one RPA70N with two peptides bound. **D**. Surface representation of the two RPA70N molecules interacting with RMI1 peptide which is displayed as olive-colored sticks.

**Fig. S6:**
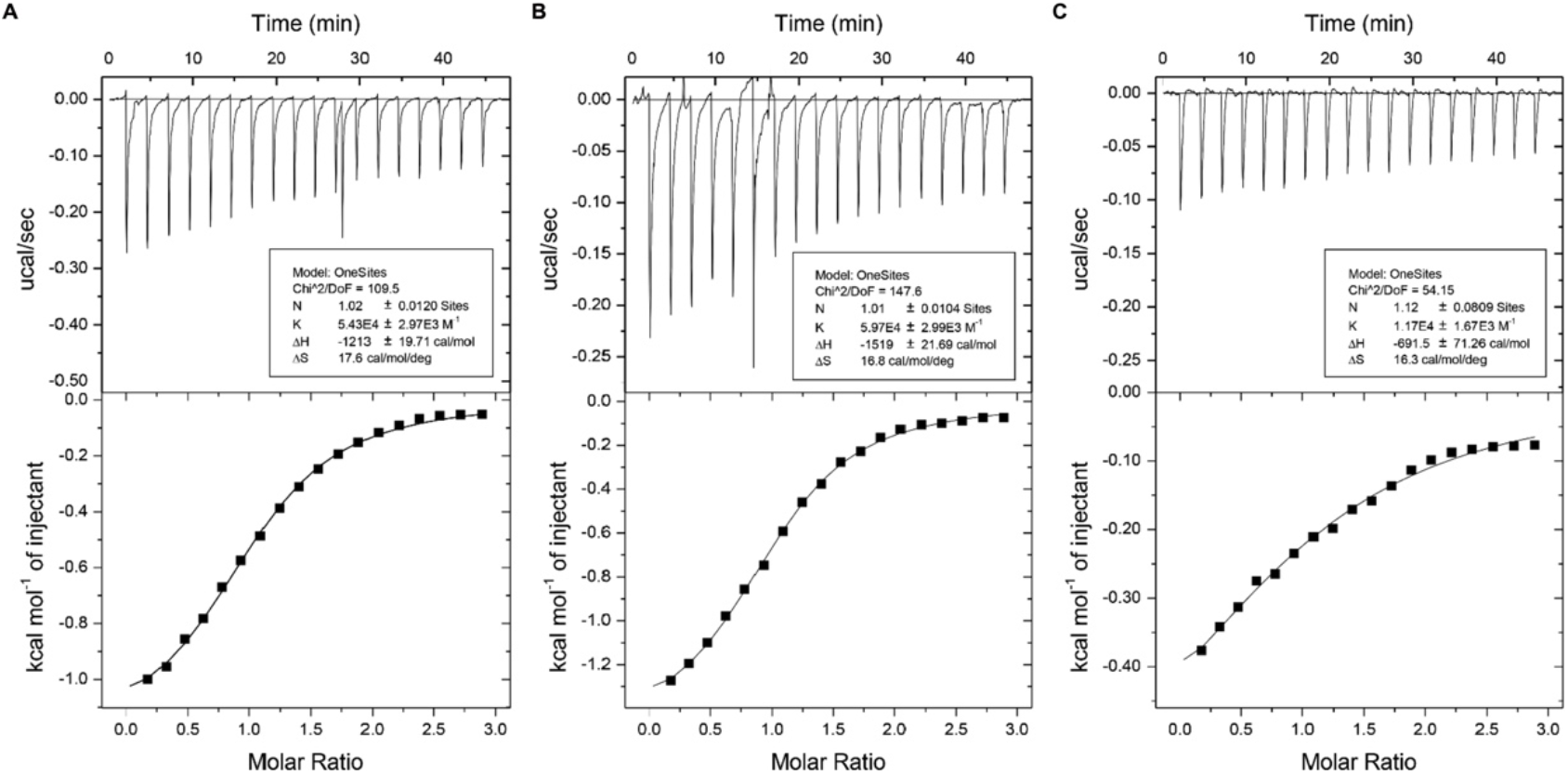
ITC titration profiles of (A) WT BLMp2 peptide (550-570aa), (B) WT BLMp1 peptide (146-165aa) or (C) BLMp1 F160A mutant peptide with RPA70N.

**Fig. S7:**
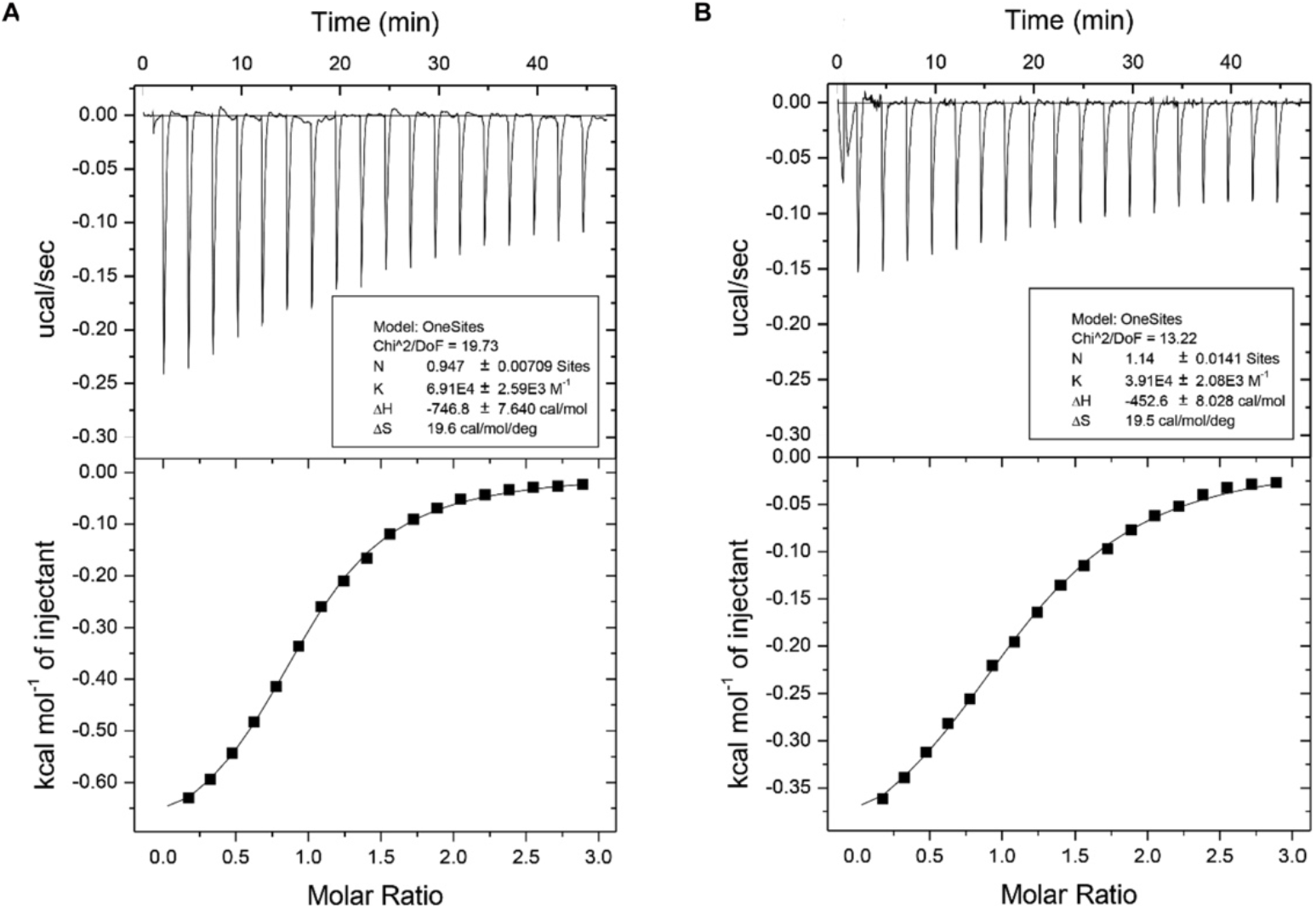
ITC titration profiles of (A) WT RMI1 peptide (243-262aa) or (B) N254A mutant peptide with RPA70N.

**Fig. S8:**
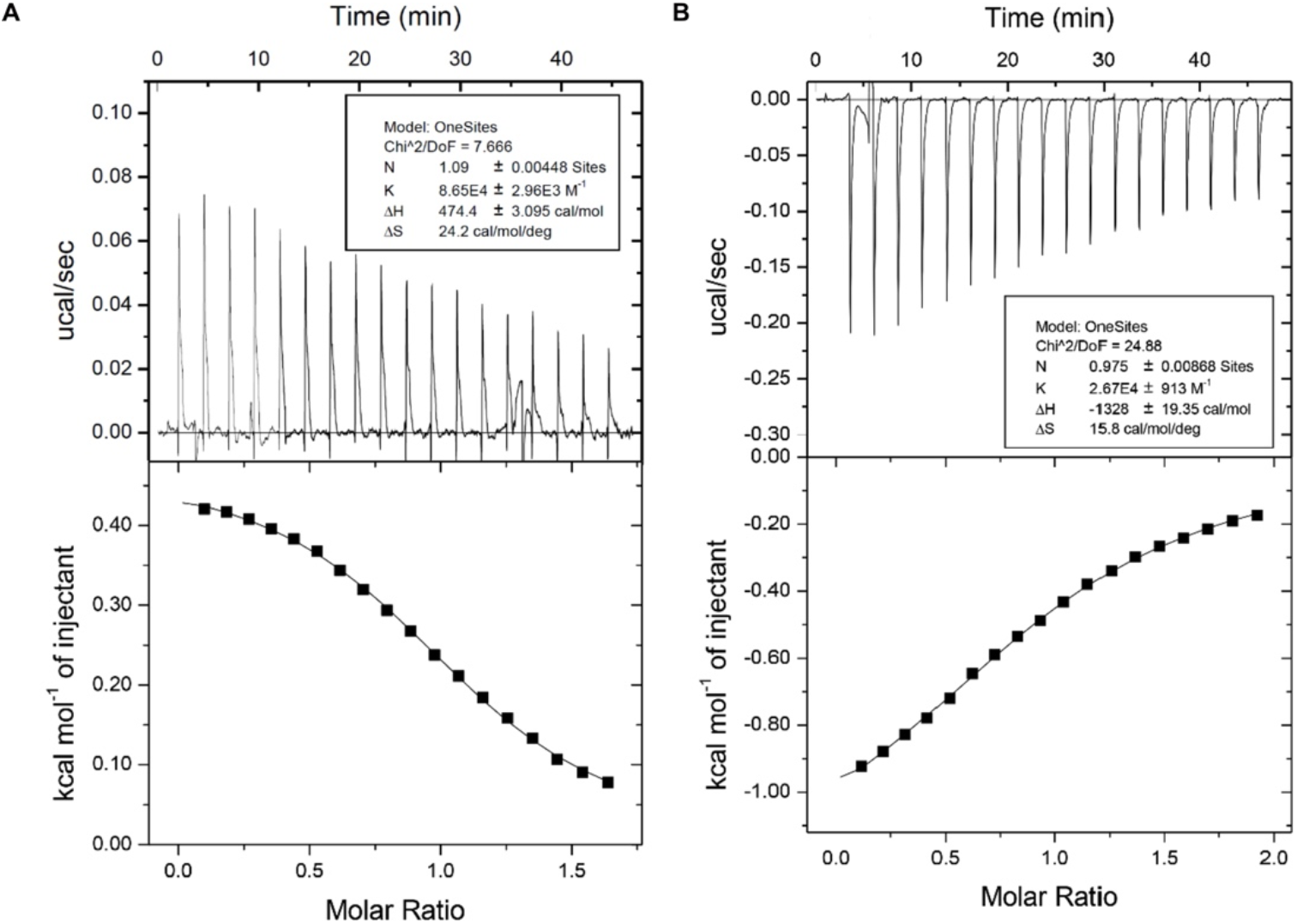
ITC titration profiles of (A) WT WRN peptide (435-451aa) or (B) WRN M466A mutant peptide with RPA70N.

**Table S1.**
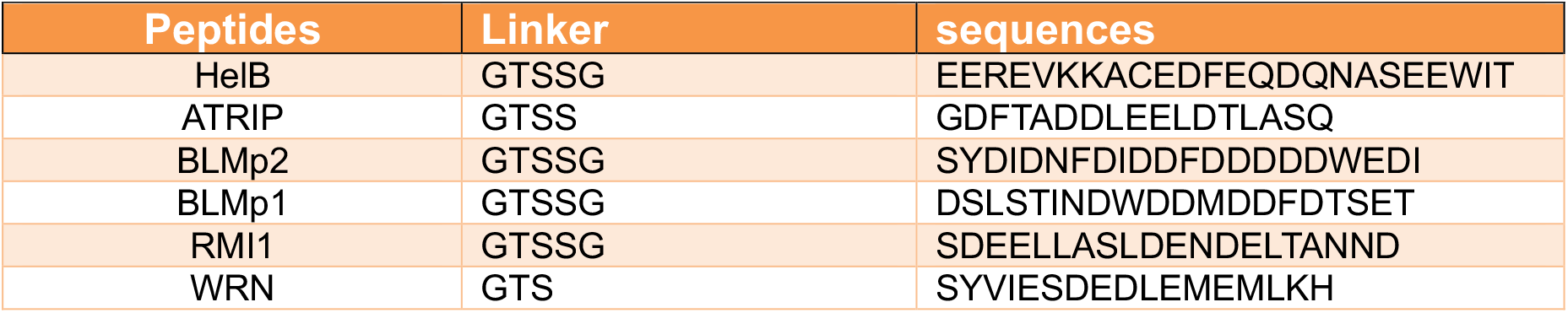
Linker and peptide sequences used in RPA70N fusion constructs.

**Table S2.**
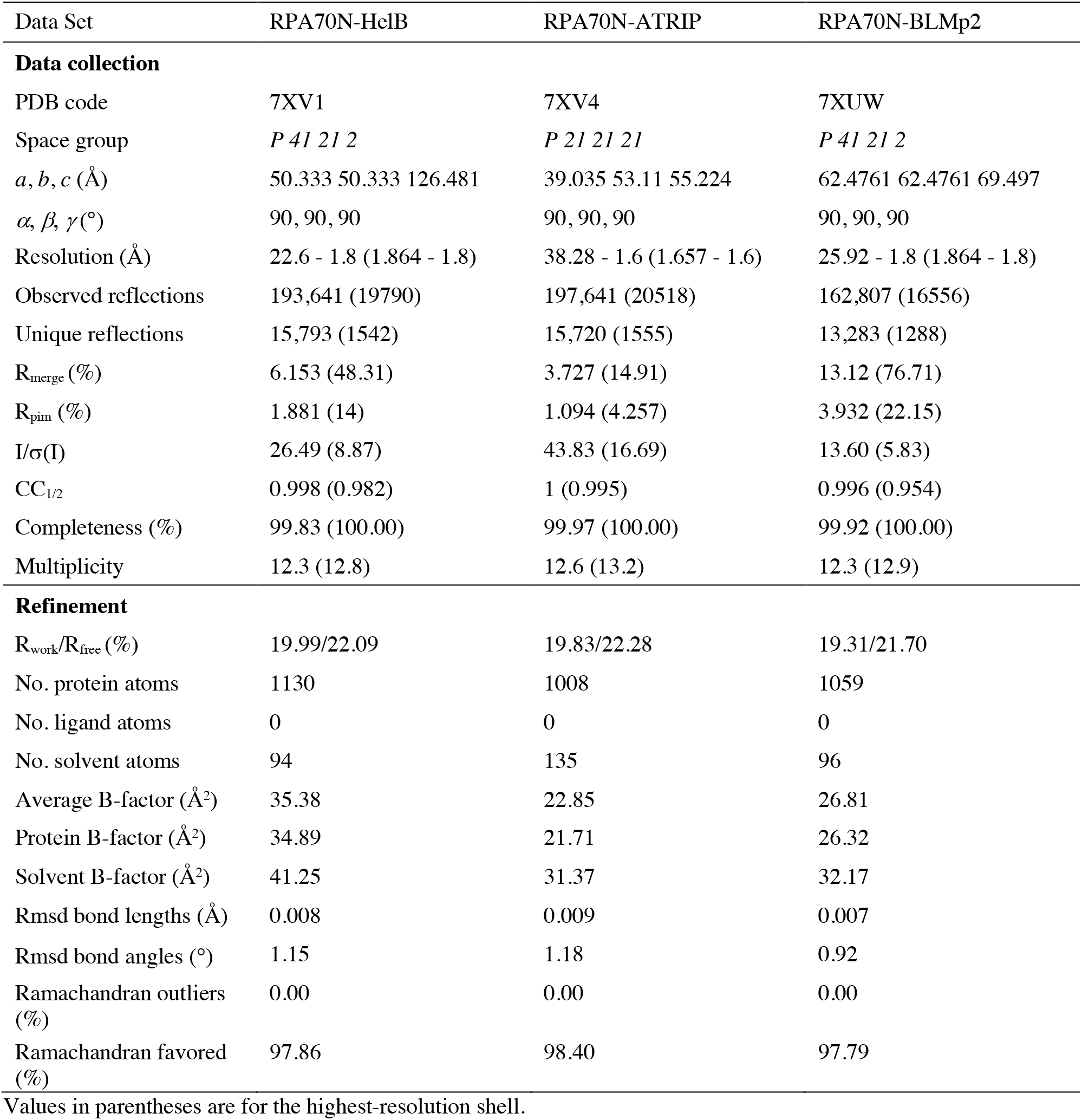
Data Collection and Refinement Statistics part 1.

**Table S3.** Data Collection and Refinement Statistics part 2.

| Data Set | RPA70N-BLMp1 | RPA70N-RMI1 | RPA70N-WRN |
| --- | --- | --- | --- |
| **Data collection** |  |  |  |
| PDB code | 7XV0 | 7XUV | 7XUT |
| Space group | *P 21 21 21* | *P 21 21 21* | *P 21 21 21* |
| *a, b, c* (Å) | 38.088 53.644 54.501 | 40.93 50.263 52.308 | 32.936 58.609 111.786 |
| $\alpha, \beta, \gamma$ (°) | 90, 90, 90 | 90, 90, 90 | 90, 90, 90 |
| Resolution (Å) | 31.22 - 1.5 (1.554 - 1.5) | 27.13 - 1.6 (1.657 - 1.6) | 31.44 - 1.6 (1.657 - 1.6) |
| Observed reflections | 111,286 (10924) | 60,739 (6230) | 181,925 (18864) |
| Unique reflections | 18,207 (1767) | 13,228 (1355) | 29,352 (2870) |
| R<sub>merge</sub> (%) | 3.1 (8.807) | 6.908 (34.22) | 9.657 (15.2) |
| R<sub>pim</sub> (%) | 1.362 (3.812) | 3.173 (15.78) | 4.187 (6.41) |
| I/σ(I) | 34.45 (16.08) | 13.80 (4.84) | 13.82 (8.55) |
| CC<sub>1/2</sub> | 0.999 (0.996) | 0.997 (0.953) | 0.992 (0.98) |
| Completeness (%) | 98.58 (98.44) | 89.41 (93.11) | 99.78 (100.00) |
| Multiplicity | 6.1 (6.2) | 4.6 (4.6) | 6.2 (6.6) |
| **Refinement** |  |  |  |
| R<sub>work</sub>/R<sub>free</sub> (%) | 16.88/20.03 | 19.08/21.88 | 18.17/21.44 |
| No. protein atoms | 1050 | 1016 | 2137 |
| No. ligand atoms | 0 | 0 | 0 |
| No. solvent atoms | 132 | 110 | 249 |
| Average B-factor (Å<sup>2</sup>) | 18.73 | 21.27 | 21.86 |
| Protein B-factor (Å<sup>2</sup>) | 17.54 | 20.36 | 21.17 |
| Solvent B-factor (Å<sup>2</sup>) | 28.19 | 29.65 | 27.76 |
| Rmsd bond lengths (Å) | 0.008 | 0.006 | 0.008 |
| Rmsd bond angles (°) | 0.92 | 1.00 | 1.02 |
| Ramachandran outliers (%) | 0.00 | 0.00 | 0.00 |
| Ramachandran favored (%) | 97.69 | 97.64 | 98.18 |

Values in parentheses are for the highest-resolution shell.

## References

Abramova NA, Russell J, Botchan M, Li R (1997) Interaction between replication protein A and p53 is disrupted after UV damage in a DNA repair-dependent manner. Proc Natl Acad Sci U S A 94: 7186–7191

Awate S, Brosh RM, Jr. (2017) Interactive Roles of DNA Helicases and Translocases with the Single-Stranded DNA Binding Protein RPA in Nucleic Acid Metabolism. Int J Mol Sci 18

Ball HL, Myers JS, Cortez D (2005) ATRIP binding to replication protein A-single-stranded DNA promotes ATR-ATRIP localization but is dispensable for Chk1 phosphorylation. Mol Biol Cell 16: 2372–2381

Binz SK, Lao Y, Lowry DF, Wold MS (2003) The phosphorylation domain of the 32-kDa subunit of replication protein A (RPA) modulates RPA-DNA interactions. Evidence for an intersubunit interaction. J Biol Chem 278: 35584–35591

Binz SK, Wold MS (2008) Regulatory functions of the N-terminal domain of the 70-kDa subunit of replication protein A (RPA). J Biol Chem 283: 21559–21570

Blackwell LJ, Borowiec JA (1994) Human replication protein A binds single-stranded DNA in two distinct complexes. Mol Cell Biol 14: 3993–4001

Bochkarev A, Pfuetzner RA, Edwards AM, Frappier L (1997) Structure of the single-stranded-DNA-binding domain of replication protein A bound to DNA. Nature 385: 176–181

Bochkareva E, Kaustov L, Ayed A, Yi GS, Lu Y, Pineda-Lucena A, Liao JC, Okorokov AL, Milner J, Arrowsmith CH et al (2005) Single-stranded DNA mimicry in the p53 transactivation domain interaction with replication protein A. Proc Natl Acad Sci U S A 102: 15412–15417

Bochkareva E, Korolev S, Lees-Miller SP, Bochkarev A (2002) Structure of the RPA trimerization core and its role in the multistep DNA-binding mechanism of RPA. EMBO J 21: 1855–1863

Brosh RM, Jr., Li JL, Kenny MK, Karow JK, Cooper MP, Kureekattil RP, Hickson ID, Bohr VA (2000) Replication protein A physically interacts with the Bloom’s syndrome protein and stimulates its helicase activity. J Biol Chem 275: 23500–23508

Brosh RM, Jr., Orren DK, Nehlin JO, Ravn PH, Kenny MK, Machwe A, Bohr VA (1999) Functional and physical interaction between WRN helicase and human replication protein A. J Biol Chem 274: 18341–18350

Bythell-Douglas R, Deans AJ (2021) A Structural Guide to the Bloom Syndrome Complex. Structure 29: 99–113

Caldwell CC, Spies M (2020) Dynamic elements of replication protein A at the crossroads of DNA replication, recombination, and repair. Crit Rev Biochem Mol Biol 55: 482–507

Cejka P, Cannavo E, Polaczek P, Masuda-Sasa T, Pokharel S, Campbell JL, Kowalczykowski SC (2010) DNA end resection by Dna2-Sgs1-RPA and its stimulation by Top3-Rmi1 and Mre11-Rad50-Xrs2. Nature 467: 112–116

Chen R, Wold MS (2014) Replication protein A: single-stranded DNA’s first responder: dynamic DNA-interactions allow replication protein A to direct single-strand DNA intermediates into different pathways for synthesis or repair. Bioessays 36: 1156–1161

Cho Y, Gorina S, Jeffrey PD, Pavletich NP (1994) Crystal structure of a p53 tumor suppressor-DNA complex: understanding tumorigenic mutations. Science 265: 346–355

Chu WK, Hickson ID (2009) RecQ helicases: multifunctional genome caretakers. Nat Rev Cancer 9: 644–654

Compton SA, Tolun G, Kamath-Loeb AS, Loeb LA, Griffith JD (2008) The Werner syndrome protein binds replication fork and holliday junction DNAs as an oligomer. J Biol Chem 283: 24478–24483

Croteau DL, Popuri V, Opresko PL, Bohr VA (2014) Human RecQ helicases in DNA repair, recombination, and replication. Annu Rev Biochem 83: 519–552

Deshpande I, Seeber A, Shimada K, Keusch JJ, Gut H, Gasser SM (2017) Structural Basis of Mec1-Ddc2-RPA Assembly and Activation on Single-Stranded DNA at Sites of Damage. Mol Cell 68: 431–445 e435

Doherty KM, Sommers JA, Gray MD, Lee JW, von Kobbe C, Thoma NH, Kureekattil RP, Kenny MK, Brosh RM, Jr. (2005) Physical and functional mapping of the replication protein a interaction domain of the werner and bloom syndrome helicases. J Biol Chem 280: 29494–29505

Dornreiter I, Erdile LF, Gilbert IU, von Winkler D, Kelly TJ, Fanning E (1992) Interaction of DNA polymerase alpha-primase with cellular replication protein A and SV40 T antigen. EMBO J 11: 769–776

Dutta A, Ruppert JM, Aster JC, Winchester E (1993) Inhibition of DNA replication factor RPA by p53. Nature 365: 79–82

Emsley P, Cowtan K (2004) Coot: model-building tools for molecular graphics. Acta Crystallogr D Biol Crystallogr 60: 2126–2132

Evans PR, Murshudov GN (2013) How good are my data and what is the resolution? Acta Crystallogr D Biol Crystallogr 69: 1204–1214

Fairman MP, Stillman B (1988) Cellular factors required for multiple stages of SV40 DNA replication in vitro. EMBO J 7: 1211–1218

Fan J, Pavletich NP (2012) Structure and conformational change of a replication protein A heterotrimer bound to ssDNA. Genes Dev 26: 2337–2347

Fanning E, Klimovich V, Nager AR (2006) A dynamic model for replication protein A (RPA) function in DNA processing pathways. Nucleic Acids Res 34: 4126–4137

Flynn RL, Zou L (2010) Oligonucleotide/oligosaccharide-binding fold proteins: a growing family of genome guardians. Crit Rev Biochem Mol Biol 45: 266–275

Frank AO, Vangamudi B, Feldkamp MD, Souza-Fagundes EM, Luzwick JW, Cortez D, Olejniczak ET, Waterson AG, Rossanese OW, Chazin WJ et al (2014) Discovery of a potent stapled helix peptide that binds to the 70N domain of replication protein A. J Med Chem 57: 2455–2461

Gravel S, Chapman JR, Magill C, Jackson SP (2008) DNA helicases Sgs1 and BLM promote DNA double-strand break resection. Genes Dev 22: 2767–2772

Guilliam TA, Brissett NC, Ehlinger A, Keen BA, Kolesar P, Taylor EM, Bailey LJ, Lindsay HD, Chazin WJ, Doherty AJ (2017) Molecular basis for PrimPol recruitment to replication forks by RPA. Nat Commun 8: 15222

Guler GD, Liu H, Vaithiyalingam S, Arnett DR, Kremmer E, Chazin WJ, Fanning E (2012) Human DNA helicase B (HDHB) binds to replication protein A and facilitates cellular recovery from replication stress. J Biol Chem 287: 6469–6481

Gupta R, Sharma S, Sommers JA, Kenny MK, Cantor SB, Brosh RM, Jr. (2007) FANCJ (BACH1) helicase forms DNA damage inducible foci with replication protein A and interacts physically and functionally with the single-stranded DNA-binding protein. Blood 110: 2390–2398

Hazeslip L, Zafar MK, Chauhan MZ, Byrd AK (2020) Genome Maintenance by DNA Helicase B. Genes (Basel) 11

Hegnauer AM, Hustedt N, Shimada K, Pike BL, Vogel M, Amsler P, Rubin SM, van Leeuwen F, Guenole A, van Attikum H et al (2012) An N-terminal acidic region of Sgs1 interacts with Rpa70 and recruits Rad53 kinase to stalled forks. EMBO J 31: 3768–3783

Hodson C, Low JKK, van Twest S, Jones SE, Swuec P, Murphy V, Tsukada K, Fawkes M, Bythell-Douglas R, Davies A et al (2022) Mechanism of Bloom syndrome complex assembly required for double Holliday junction dissolution and genome stability. Proc Natl Acad Sci U S A 119

Hormeno S, Wilkinson OJ, Aicart-Ramos C, Kuppa S, Antony E, Dillingham MS, Moreno-Herrero F (2022) Human HELB is a processive motor protein that catalyzes RPA clearance from single-stranded DNA. Proc Natl Acad Sci U S A 119: e2112376119

Iftode C, Daniely Y, Borowiec JA (1999) Replication protein A (RPA): the eukaryotic SSB. Crit Rev Biochem Mol Biol 34: 141–180

Jacobs DM, Lipton AS, Isern NG, Daughdrill GW, Lowry DF, Gomes X, Wold MS (1999) Human replication protein A: global fold of the N-terminal RPA-70 domain reveals a basic cleft and flexible C-terminal linker. J Biomol NMR 14: 321–331

Kabsch W (2010) Xds. Acta Crystallogr D Biol Crystallogr 66: 125–132

Kang D, Lee S, Ryu KS, Cheong HK, Kim EH, Park CJ (2018) Interaction of replication protein A with two acidic peptides from human Bloom syndrome protein. FEBS Lett 592: 547–558

Kim C, Paulus BF, Wold MS (1994) Interactions of human replication protein A with oligonucleotides. Biochemistry 33: 14197–14206

Kim C, Snyder RO, Wold MS (1992) Binding properties of replication protein A from human and yeast cells. Mol Cell Biol 12: 3050–3059

Kitano K (2014) Structural mechanisms of human RecQ helicases WRN and BLM. Front Genet 5: 366

Lee M, Shin S, Uhm H, Hong H, Kirk J, Hyun K, Kulikowicz T, Kim J, Ahn B, Bohr VA et al (2018) Multiple RPAs make WRN syndrome protein a superhelicase. Nucleic Acids Res 46: 4689–4698

Lee S, Heo J, Park CJ (2020) Determinants of replication protein A subunit interactions revealed using a phosphomimetic peptide. J Biol Chem 295: 18449–18458

Li R, Botchan MR (1993) The acidic transcriptional activation domains of VP16 and p53 bind the cellular replication protein A and stimulate in vitro BPV-1 DNA replication. Cell 73: 1207–1221

Liebschner D, Afonine PV, Baker ML, Bunkoczi G, Chen VB, Croll TI, Hintze B, Hung LW, Jain S, McCoy AJ et al (2019) Macromolecular structure determination using X-rays, neutrons and electrons: recent developments in Phenix. Acta Crystallogr D Struct Biol 75: 861–877

Lindsey-Boltz LA, Reardon JT, Wold MS, Sancar A (2012) In vitro analysis of the role of replication protein A (RPA) and RPA phosphorylation in ATR-mediated checkpoint signaling. J Biol Chem 287: 36123–36131

Liu Y, Vaithiyalingam S, Shi Q, Chazin WJ, Zinkel SS (2011) BID binds to replication protein A and stimulates ATR function following replicative stress. Mol Cell Biol 31: 4298–4309

Marechal A, Zou L (2015) RPA-coated single-stranded DNA as a platform for post-translational modifications in the DNA damage response. Cell Res 25: 9–23

McCoy AJ, Grosse-Kunstleve RW, Adams PD, Winn MD, Storoni LC, Read RJ (2007) Phaser crystallographic software. J Appl Crystallogr 40: 658–674

Mukherjee S, Sinha D, Bhattacharya S, Srinivasan K, Abdisalaam S, Asaithamby A (2018) Werner Syndrome Protein and DNA Replication. Int J Mol Sci 19

Murzin AG (1993) OB(oligonucleotide/oligosaccharide binding)-fold: common structural and functional solution for non-homologous sequences. EMBO J 12: 861–867

Namiki Y, Zou L (2006) ATRIP associates with replication protein A-coated ssDNA through multiple interactions. Proc Natl Acad Sci U S A 103: 580–585

Nimonkar AV, Genschel J, Kinoshita E, Polaczek P, Campbell JL, Wyman C, Modrich P, Kowalczykowski SC (2011) BLM-DNA2-RPA-MRN and EXO1-BLM-RPA-MRN constitute two DNA end resection machineries for human DNA break repair. Genes Dev 25: 350–362

Ning B, Feldkamp MD, Cortez D, Chazin WJ, Friedman KL, Fanning E (2015) Simian virus Large T antigen interacts with the N-terminal domain of the 70 kD subunit of Replication Protein A in the same mode as multiple DNA damage response factors. PLoS One 10: e0116093

Oakley GG, Tillison K, Opiyo SA, Glanzer JG, Horn JM, Patrick SM (2009) Physical interaction between replication protein A (RPA) and MRN: involvement of RPA2 phosphorylation and the N-terminus of RPA1. Biochemistry 48: 7473–7481

Robison JG, Elliott J, Dixon K, Oakley GG (2004) Replication protein A and the Mre11.Rad50.Nbs1 complex co-localize and interact at sites of stalled replication forks. J Biol Chem 279: 34802–34810

Shen JC, Gray MD, Oshima J, Loeb LA (1998) Characterization of Werner syndrome protein DNA helicase activity: directionality, substrate dependence and stimulation by replication protein A. Nucleic Acids Res 26: 2879–2885

Shen JC, Lao Y, Kamath-Loeb A, Wold MS, Loeb LA (2003) The N-terminal domain of the large subunit of human replication protein A binds to Werner syndrome protein and stimulates helicase activity. Mech Ageing Dev 124: 921–930

Shorrocks AK, Jones SE, Tsukada K, Morrow CA, Belblidia Z, Shen J, Vendrell I, Fischer R, Kessler BM, Blackford AN (2021) The Bloom syndrome complex senses RPA-coated single-stranded DNA to restart stalled replication forks. Nat Commun 12: 585

Soniat MM, Myler LR, Kuo HC, Paull TT, Finkelstein IJ (2019) RPA Phosphorylation Inhibits DNA Resection. Mol Cell 75: 145–153 e145

Souza-Fagundes EM, Frank AO, Feldkamp MD, Dorset DC, Chazin WJ, Rossanese OW, Olejniczak ET, Fesik SW (2012) A high-throughput fluorescence polarization anisotropy assay for the 70N domain of replication protein A. Anal Biochem 421: 742–749

Suhasini AN, Sommers JA, Mason AC, Voloshin ON, Camerini-Otero RD, Wold MS, Brosh RM, Jr. (2009) FANCJ helicase uniquely senses oxidative base damage in either strand of duplex DNA and is stimulated by replication protein A to unwind the damaged DNA substrate in a strand-specific manner. J Biol Chem 284: 18458–18470

Taneja P, Gu J, Peng R, Carrick R, Uchiumi F, Ott RD, Gustafson E, Podust VN, Fanning E (2002) A dominant-negative mutant of human DNA helicase B blocks the onset of chromosomal DNA replication. J Biol Chem 277: 40853–40861

Tkac J, Xu G, Adhikary H, Young JTF, Gallo D, Escribano-Diaz C, Krietsch J, Orthwein A, Munro M, Sol W et al (2016) HELB Is a Feedback Inhibitor of DNA End Resection. Mol Cell 61: 405–418

Wan L, Lou J, Xia Y, Su B, Liu T, Cui J, Sun Y, Lou H, Huang J (2013) hPrimpol1/CCDC111 is a human DNA primase-polymerase required for the maintenance of genome integrity. EMBO Rep 14: 1104–1112

Wang F, Yang Y, Singh TR, Busygina V, Guo R, Wan K, Wang W, Sung P, Meetei AR, Lei M (2010) Crystal structures of RMI1 and RMI2, two OB-fold regulatory subunits of the BLM complex. Structure 18: 1159–1170

Winn MD, Ballard CC, Cowtan KD, Dodson EJ, Emsley P, Evans PR, Keegan RM, Krissinel EB, Leslie AG, McCoy A et al (2011) Overview of the CCP4 suite and current developments. Acta Crystallogr D Biol Crystallogr 67: 235–242

Wold MS (1997) Replication protein A: a heterotrimeric, single-stranded DNA-binding protein required for eukaryotic DNA metabolism. Annu Rev Biochem 66: 61–92

Wood RD, Robins P, Lindahl T (1988) Complementation of the xeroderma pigmentosum DNA repair defect in cell-free extracts. Cell 53: 97–106

Wu X, Shell SM, Zou Y (2005) Interaction and colocalization of Rad9/Rad1/Hus1 checkpoint complex with replication protein A in human cells. Oncogene 24: 4728–4735

Xu D, Guo R, Sobeck A, Bachrati CZ, Yang J, Enomoto T, Brown GW, Hoatlin ME, Hickson ID, Wang W (2008a) RMI, a new OB-fold complex essential for Bloom syndrome protein to maintain genome stability. Genes Dev 22: 2843–2855

Xu X, Vaithiyalingam S, Glick GG, Mordes DA, Chazin WJ, Cortez D (2008b) The basic cleft of RPA70N binds multiple checkpoint proteins, including RAD9, to regulate ATR signaling. Mol Cell Biol 28: 7345–7353

Xue X, Raynard S, Busygina V, Singh AK, Sung P (2013) Role of replication protein A in double holliday junction dissolution mediated by the BLM-Topo IIIalpha-RMI1-RMI2 protein complex. J Biol Chem 288: 14221–14227

Yates LA, Aramayo RJ, Pokhrel N, Caldwell CC, Kaplan JA, Perera RL, Spies M, Antony E, Zhang X (2018) A structural and dynamic model for the assembly of Replication Protein A on single-stranded DNA. Nat Commun 9: 5447

Yeom G, Kim J, Park CJ (2019) Investigation of the core binding regions of human Werner syndrome and Fanconi anemia group J helicases on replication protein A. Sci Rep 9: 14016

Zhou C, Pourmal S, Pavletich NP (2015) Dna2 nuclease-helicase structure, mechanism and regulation by Rpa. Elife 4

Zhu Z, Chung WH, Shim EY, Lee SE, Ira G (2008) Sgs1 helicase and two nucleases Dna2 and Exo1 resect DNA double-strand break ends. Cell 134: 981–994

Zou L, Elledge SJ (2003) Sensing DNA damage through ATRIP recognition of RPA-ssDNA complexes. Science 300: 1542–1548

Zou Y, Liu Y, Wu X, Shell SM (2006) Functions of human replication protein A (RPA): from DNA replication to DNA damage and stress responses. J Cell Physiol 208: 267–273

